# IceR improves proteome coverage and data completeness in global and single-cell proteomics

**DOI:** 10.1101/2020.11.01.363101

**Authors:** Mathias Kalxdorf, Torsten Müller, Oliver Stegle, Jeroen Krijgsveld

## Abstract

Label-free proteomics by data-dependent acquisition (DDA) enables the unbiased quantification of thousands of proteins, however it notoriously suffers from high rates of missing values, thus prohibiting consistent protein quantification across large sample cohorts. To solve this, we here present IceR, an efficient and user-friendly quantification workflow that combines high identification rates of DDA with low missing value rates similar to DIA. Specifically, IceR uses ion current information in DDA data for a hybrid peptide identification propagation (PIP) approach with superior quantification precision, accuracy, reliability and data completeness compared to other quantitative workflows. We demonstrate greatly improved quantification sensitivity on published plasma and single-cell proteomics data, enhancing the number of reliably quantified proteins, improving discriminability between single-cell populations, and allowing reconstruction of a developmental trajectory. IceR will be useful to improve performance of large scale global as well as low-input proteomics applications, facilitated by its availability as an easy-to-use R-package.

## Introduction

The reproducible quantification of peptides and proteins across large sample cohorts is crucial to investigate proteome differences between individuals and across conditions^1^. Mass spectrometry is the leading technology to achieve this, operating to identify and quantify peptides that are generated from cell or tissue lysates, often in conjunction with liquid chromatography to maximize the number of detected peptides and thus enhance sampling depth^2^. The two main experimental approaches operate either via data-dependent or data-independent acquisition (DDA and DIA, respectively)^3^. In the traditional and more commonly used DDA approach, a pre-defined number of most abundant precursor ions (‘Top N’) detected in a MS1 survey scan are sequentially selected for MS2-based peptide fragmentation and protein identification^4^. Although this has been successfully used to characterize thousands of proteins in countless proteomic studies, a large fraction of peptide ions detected in MS1 is typically not targeted for fragmentation due to stochasticity of precursor selection, thus leading to missing data^5,6^. This is a persistent problem caused by proteome complexity and dynamic range that cannot be fully captured, despite continuous improvements of sensitivity and acquisition speeds of mass spectrometers^7^. The degree of missing data becomes increasingly prominent in complex samples and in large sample cohorts^8^, resulting in decreased numbers of proteins that can be consistently identified across all specimens^9^, and hence limiting the power of DDA in large-scale proteomics e.g. for biomarker discovery. To circumvent this, several experimental and computational strategies have been developed in the last years to overcome the problem of missing values in DDA. First, DIA approaches have been introduced, co-fragmenting peptides in bins across the entire peptide mass range, resulting in considerable advancements in reproducible protein measurement^10,11^. However, DIA suffers from limited depth of quantifications due to the burden of peptide identification from severely convoluted data^8,12^, and it requires an upfront reference library typically acquired by DDA^13^. Because of these limitations of DIA and because of historical reasons, DDA is still the most frequently applied approach in label-free proteomics. This also applies to recent efforts to advance proteomics methods for low input regimes, including single-cell analyses^14^. Second, multiplexed labeling strategies such as TMT have been introduced, allowing the simultaneous detection and quantification of peptides in an 11 or even 16-plexed manner at significantly reduced rates of missing values^15^. This approach has also been used for single-cell analysis, using one of the TMT channels to boost the signal in MS1 to benefit detection of peptides in the other channels^14,16^. Yet, the optimal size of the booster channel as well as quantification of low-abundance features in the single-cell channels remain subject of debate^17^. Third, missing values can be replaced post hoc by various imputation strategies, which can be highly challenging since peptides may be missing for various reasons, even within individual samples^18–20^. Finally, peptide identity propagation (PIP) provides a powerful approach to transfer sequencing information between samples, enabling the assignment of a peptide identifier to a feature even if it has not been selected for fragmentation^8^. This can be performed by feature-based or ion-based PIP, both requiring accurate retention time and m/z alignment over samples. Feature-based PIP additionally requires the presence of molecular ions to be observed as isotope peak patterns^21^, which can be performed with traditional feature detection algorithms as has been implemented by the match-between-runs (MBR) algorithm in the MaxQuant environment^21,22^. However, the need to recognize isotope peak patterns limits the sensitivity of this approach, thereby preventing feature-based PIP from fully solving the missing-value problem^8^. In addition, false transfers cannot be excluded, requiring specific attention^23^. In contrast, ion-based PIP applies direct ion current extraction (DICE), and only requires the existence of ions within a given retention time and mass-to-charge window, thus enabling sensitive identity propagation^1^. Ion-based PIP has been implemented in DeMix-Q^8^ and IonStar^12^, both achieving highly reduced missing value rates and improved sensitivity to detect differentially abundant proteins compared to MaxQuant. However, without rigorous quality control measures, this goes at the cost of lower matching reliability^8^ as DICE is less constrained. Further, both tools suffer from several issues that prevent their general use: For instance, they apply fixed and large ion current extraction windows (e.g. m/z ± 5 ppm, RT ± 1 min) resulting in data deterioration by co-eluting interferences; they do not distinguish between true presence of a peptide and random occurrence of an ion within the quantification windows; they are designed to only process data from Thermo Fisher Scientific Orbitrap mass spectrometers, limiting their use to other vendors and scan modes (e.g. ion mobility (IM) separation). In addition, running of DeMix-Q and IonStar is highly cumbersome, requiring installation of several tools including discontinued commercial applications. This may explain why DICE-based approaches, despite their advantages, have never permeated into mainstream proteomics applications.

It is easy to argue why the principle of DICE is advantageous over other PIP methods, allowing sensitive feature detection and making post hoc imputation obsolete, however its implementation, especially in a user-friendly manner, has proven not straightforward. Therefore, we here present IceR (Ion current extraction Re-quantification), an efficient, robust, and user-friendly label-free proteomics quantification workflow. The method uniquely combines the following features: 1) a hybrid PIP approach merging feature-based and ion-based PIP; 2) robust 2-step feature alignment incorporating global modelling-based and local feature-specific kernel density estimation, allowing narrow and feature-specific DICE-windows in m/z-, RT- and IM-space; 3) capability to utilize ion mobility as an additional dimension for feature detection and alignment; 4) sound decoy feature-based scoring schemes to assess reliability of quantifications and to distinguish true presence of peptides from random ion occurrences; 5) a superior noise-model-based imputation approach allowing accurate estimations of ratios of low abundance peptides and proteins. Owing to the combination of these features, IceR enables superior quantification precision, accuracy, reliability, and data completeness. To allow for broad and easy applicability, we have implemented IceR as a user-friendly R-package (will be made publicly available on GitHub). It can be seamlessly integrated with the MaxQuant suite, or with any other pre-processed label-free proteomics data set for which detected features are reported. The software provides a graphical user interface to set up analyses and inspect quality control measures.

We have comprehensively assessed and benchmarked IceR on 4 publicly available and 3 in-house generated data sets including comparisons against DeMix-Q, IonStar, DIA and IM-enhanced MaxQuant. Furthermore, we have evaluated its performance on published plasma and single-cell proteomics data sets enabling highly increased numbers of reliably quantified proteins, improving discriminability between single-cell populations and enhancing de novo reconstruction of a developmental trajectory.

## Results

### Analysis pipeline

We designed IceR to leverage DICE for peptide quantification, and to use it for reliable and sensitive PIP to minimize missing values across proteomic data sets. In addition, since DICE will primarily rescue low-intensity peptides that escaped direct identification by MS2, we aimed to assess the overall gain in sensitivity that could be achieved compared to commonly used approaches for PIP. The IceR approach is schematically summarized in Fig. 1a, and the complete workflow is described in Supplementary Text 1, and illustrated in Supplementary Fig. 1. In brief, IceR starts from lists of peptide features detected in label-free DDA data. These features are aggregated and aligned over all samples (steps 1-3 in Supplementary Fig. 1), and finally the respective quantities are extracted by DICE from MS raw files. To enable reliable PIP, IceR performs modelling-based RT- and m/z-corrections (step 4), similar to MaxQuant, DeMix-Q and IonStar. While this approach can greatly decrease global variability between samples, local sample- and feature-specific heterogeneities might be missed. To account for this, IceR uniquely performs a second sample- and feature-specific alignment step based on kernel density estimated ion accumulation maps (steps 6-8). This enables highly robust alignments over samples, and hence allows more narrow DICE windows. Furthermore, IceR incorporates a hybrid PIP approach: sequencing information is propagated preferably by feature-based PIP, and quantifications are recovered by ion-based PIP only in cases of missing feature detection (step 5). Decoy feature-based scoring schemes are applied to assess reliability of PIPs and quantifications (steps 6, 9-11). Along with a detailed description of all steps of the IceR workflow, supplementary Text 1 and supplementary Figs. 2-3 illustrate its application to a published spike-in data set (iPRG 2015 study^24^) and assesses its performance compared to MaxQuant and DeMix-Q. Among the most salient findings, i) IceR could transfer sequence information for nearly twice the number features due to its hybrid PIP approach (supplementary Fig. 2a) and its superior two-step alignment algorithm (Supplementary Fig. 2b,c); ii) FDR of DICE-based peak selection was estimated at 0.6% (Supplementary Fig. 2d); iii) on average 85% of all peptide features could be directly quantified by IceR (52% by MaxQuant) (Supplementary Fig. 2a,e); iv) IceR reduced missing value rates by 12-fold compared to MaxQuant, being at par with DeMix-Q (Supplementary Fig. 2f,g); v) IceR resulted in most accurate and precise protein abundance ratio estimations (Supplementary Fig. 2h) and enhanced statistical power for DE analyses compared to DeMix-Q (Supplementary Fig. 2i). The results of this initial analysis indicated highly promising performance of IceR, prompting us to evaluate it in an additional set of use cases.

**Fig. 1.**
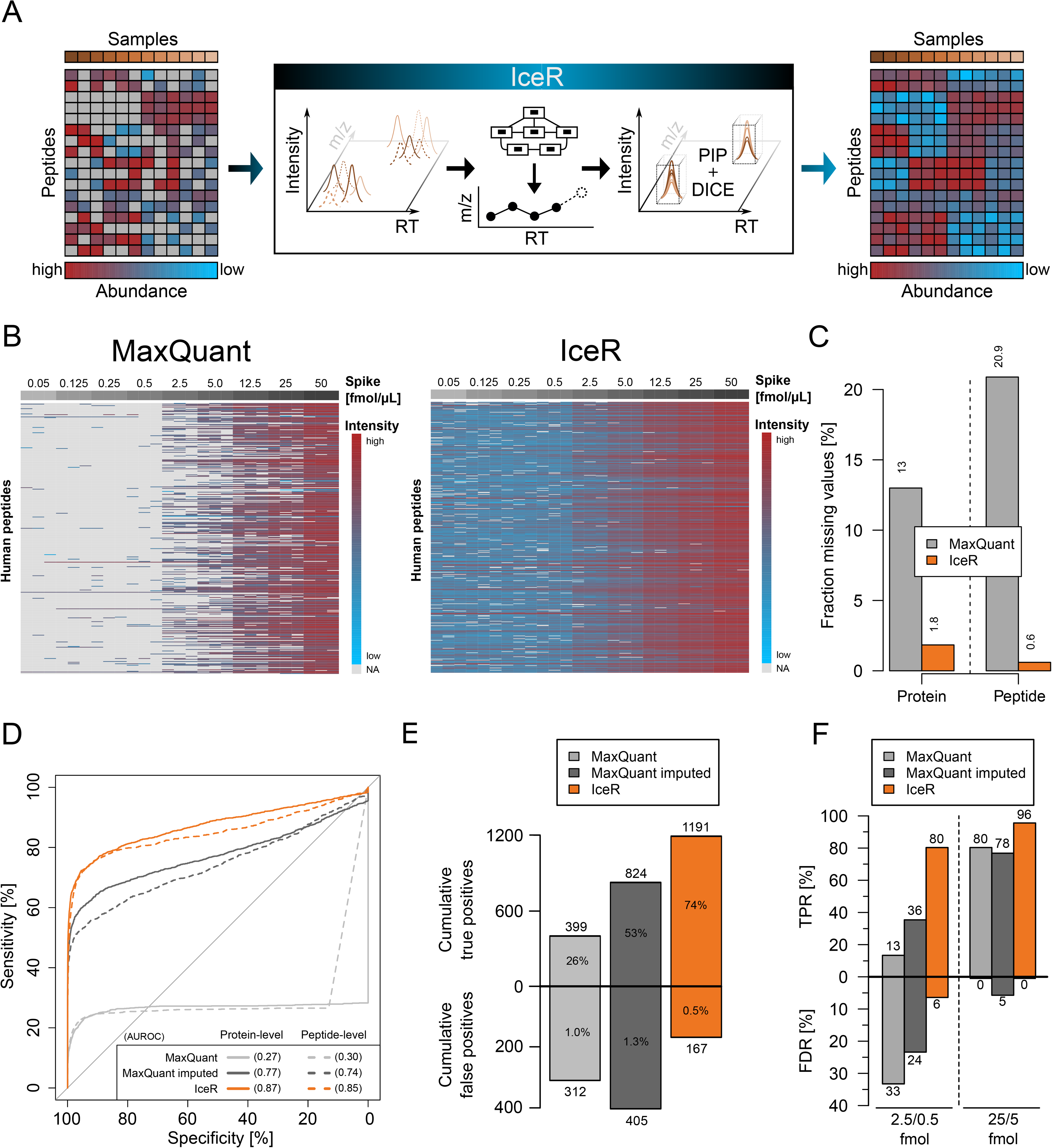
IceR enables enhanced sensitivity to detect differentially abundant proteins. **a**, Label-free DDA proteomics typically results in increasing numbers of missing values with increasing sample sizes due to the stochastic principle of precursor selection for sequencing. IceR addresses this issue by performing robust peptide feature alignment of samples in m/z and chromatographic retention time (RT) space enabling reliable peptide identity propagation (PIP) across samples and highly sensitive and accurate quantification by direct ion current extraction (DICE) quantification. Thereby, IceR highly reduces missing value rates and enables comprehensive, precise and accurate label-free proteomics analyses. **b**, Heatmap representation of quantified peptides of 48 spiked proteins at nine spike amounts into constant background (n=3) in MaxQuant (left) and IceR (right) results. Low abundant peptides are coloured blue, high abundant peptides are coloured red, and missing values are indicated in grey. **c**, Fraction of missing values on protein- and peptide-level in MaxQuant (grey) and IceR (orange) results. **d**, Receiver operating characteristics (ROC) over all pairwise differential expression analyses for MaxQuant (light grey), MaxQuant with imputation (dark grey) and IceR (orange) on protein-level (solid line) and peptide-level (dashed line). Area under the ROC (AUROC) per condition is indicated. **e**, Cumulative true and false positives over all (36) pairwise DE analyses for MaxQuant (grey), MaxQuant with imputation (dark grey) and IceR (orange). True and false positive rates are indicated. **f**, True positive rate (TPR) over false discovery rate (FDR) for differential abundance testing with a true 5-fold spike-in ratio at low (2.5 vs 0.5 fmol/μL) and high (25/5 fmol/μL) spike-in amounts in MaxQuant (light grey), MaxQuant with imputation (dark grey) and IceR (orange).

### Evaluation of IceR based on public spike-in data sets

Beyond testing against the iPRG2015 study^24^ (Supplementary Fig. 2-3), we benchmarked IceR against two additional published data sets with increasing complexity.

In the data set published by Ramus et al.^25^, 48 recombinant human proteins (UPS1 mix) were spiked into yeast lysate at 9 different concentrations ranging from 0.05 fmol/μL to 50 fmol/μL, to test the recovery of a defined set of proteins against a complex background. In total, 883 yeast proteins and 43 UPS1 proteins were identified by MaxQuant (MBR enabled) with at least 2 peptides (Supplementary Fig. 4a). When re-analysing these data with IceR, the fraction of missing values on protein-level could be reduced from 13.0% to 1.8% and on peptide-level from 20.9% to 0.6% (Fig. 1b,c). This enabled almost full quantification of the 45 identified UPS1 proteins in IceR down to the most diluted condition. Coefficients of variation (CV) of quantifications were comparably low between MaxQuant and IceR data (both 4.9% on protein-level, supplementary Fig. 4b,c), despite the fact that more than 2-times more quantification events on peptide-level were available in IceR data (48K vs 117K, Supplementary Fig. 4c), the majority of which were of low abundance (Fig. 1b). The receiver operating characteristic (ROC) showed superior performance of IceR over all pairwise DE analyses on protein- and on peptide-level (Fig. 1d). Accordingly, IceR resulted in the highest true positive rate (TPR) and lowest false positive rate (FPR) in comparison to MaxQuant with and without missing value imputation (Fig. 1e). Protein abundance ratios could be accurately estimated for almost 3.5-times more spiked proteins over all pairwise DE analyses, compared to MaxQuant data (Supplementary Fig. 4d). Furthermore, due to its ability to robustly quantify peptides and proteins even at low abundances, IceR enabled superior TPR and low false discovery rates (FDR) even at low absolute spike-in amounts (Fig. 1f and Supplementary Fig. 4e,f).

In the data set published by Shen et al.^12^, a complete *E. coli* lysate was spiked into human lysate at 5 ratios (*E. coli*/human), ranging from 3 % to 9 %. As this data set was originally used to evaluate the performance of IonStar in recovering low abundance proteins at a proteomic scale, we used it here for its direct comparison to MaxQuant and IceR. Numbers of identified proteins were comparable between MaxQuant and IceR outputs, however, IonStar identified >10% fewer proteins (Supplementary Fig. 4g). Missing values on protein-level amounted to 14.1% in MaxQuant, which was reduced to 0.1% by IceR and 0% in IonStar, thus allowing an (almost) complete quantification of identified *E. coli* proteins by the latter two approaches (Fig. 2a,b). Analysis of quantification variability revealed a comparable CV between MaxQuant and IceR data (both 6.5%) while IonStar resulted in a more variable quantification (10.1%) between replicates (Supplementary Fig. 4h,i). Still, IonStar outperformed MaxQuant in detecting differentially abundant spiked proteins in pairwise differential expression analyses as evidenced from the area under the ROC (AUROC) that was increased from 0.72 to 0.86 on protein-level (Fig. 2c). Imputing missing values in MaxQuant data abolished this AUROC difference. In contrast, IceR resulted in an almost perfect sensitivity over all pairwise differential expression analyses, indicated by an AUROC of 0.99 on protein- and peptide-level (Fig 2c). Accordingly, IceR resulted in highest TPR and lowest FPR over all pairwise DE analyses (Fig. 2d), and especially in case of small ratios (abundance difference of ≤ 50%) IceR showed superior sensitivity compared to IonStar and MaxQuant for detecting true positives at low FDR irrespective of the absolute spike-in amounts (Fig. 2e). Protein abundance ratios could be accurately estimated in all cases (Supplementary Fig. 4j). In conclusion, IceR clearly outperformed MaxQuant and IonStar by enabling low missing value rates with highly accurate and precise quantifications resulting in the highest TPR and lowest FDR.

**Fig. 2.**
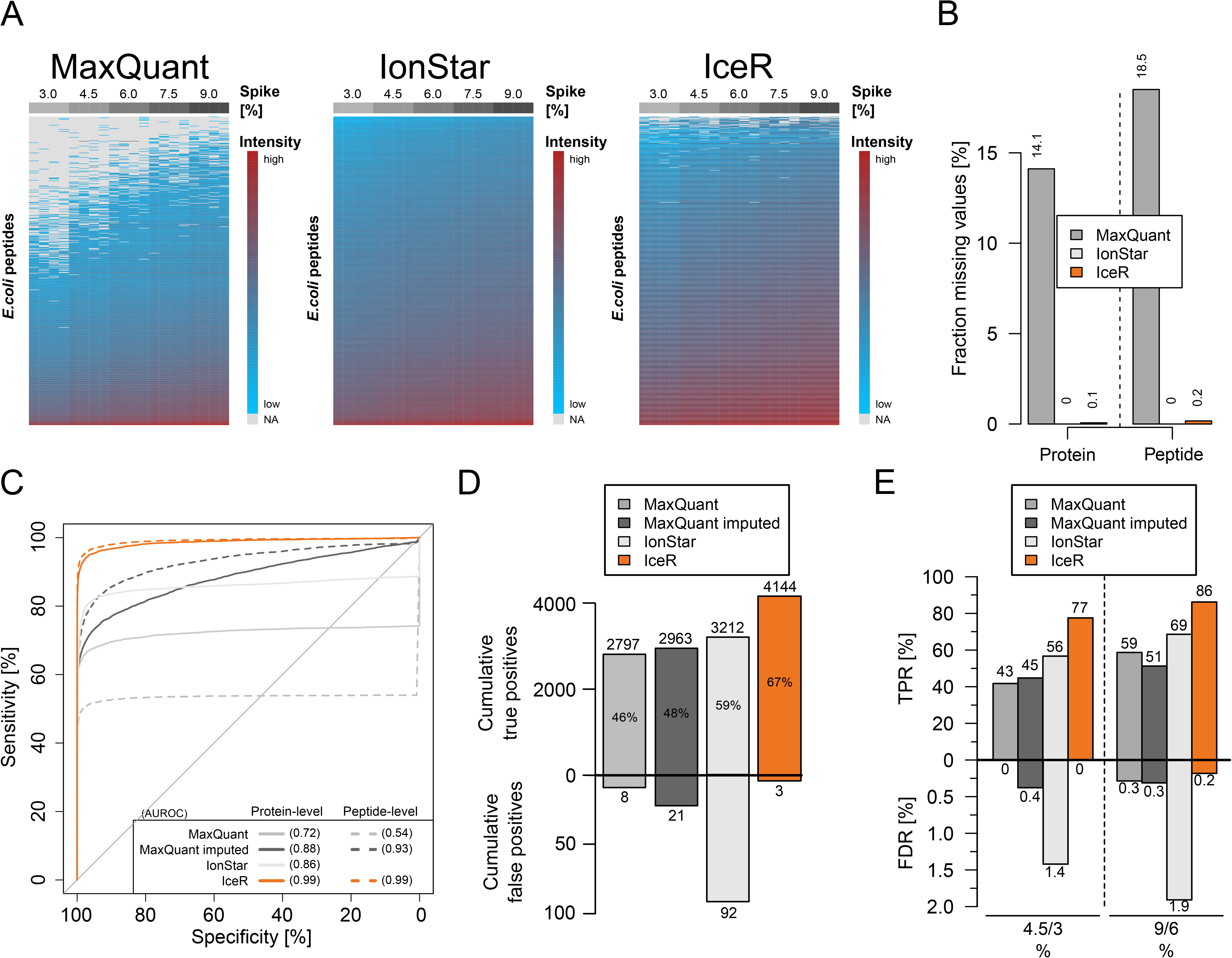
IceR outperforms other label-free quantification workflows. **a**, Heatmap representation of quantified peptides of spiked *E. coli* lysate at five spike amounts into constant background (n=4) in MaxQuant (left), IonStar (middle) and IceR (right) results. Low abundance peptides are coloured blue, high abundance peptides are coloured red, and missing values are indicated in grey. **b**, Fraction of missing values on protein- and peptide-level in MaxQuant (grey), IonStar (red) and IceR (orange) results. **c**, Receiver operating characteristics (ROC) over all pairwise differential expression analyses for MaxQuant (light grey), MaxQuant with imputation (dark grey), IonStar (red) and IceR (orange) on protein-level (solid line) and peptide-level (dashed line). Area under the ROC (AUROC) per condition is indicated. **d**, Cumulative true and false positives over all (10) pairwise DE analyses for MaxQuant (grey), MaxQuant with imputation (dark grey), IonStar (light grey) and IceR (orange). True positive rates are indicated. **e**, True positive rate (TPR) over false discovery rate (FDR) for differential abundance testing with a true 1.5-fold spike-in ratio at low (4.5 vs 3 % *E. coli*) and high (9 vs 6 % *E. coli*) spike-in amounts in MaxQuant (light grey), MaxQuant with imputation (dark grey), IonStar (red) and IceR (orange).

### Evaluation of IceR performance at various MS-acquisition conditions

Experimental parameters in MS are typically varied to maximize the number of identified and quantified proteins, however this may inflate missing values or deteriorate reliability of quantification. To evaluate how IceR performs within typical parameter ranges, an *E. coli* lysate was spiked into constant human lysate in 6 different amounts (0, 3, 4.5, 6, 7.5 and 9% relative to human, wt/wt, n=3). Respective samples were analysed by changing one parameter while keeping the other parameters constant. We tested: 1) ‘Top N’ (Top5, Top10 and Top20); 2) Gradient length (1h, 1h25 and 2h); and 3) Sample amount (500 ng, 50 ng, and 10 ng).

In all tested scenarios, IceR outperformed MaxQuant results: It enabled identification of more *E. coli* proteins with at least 2 quantification events (Fig. 3a, top panel), resulted in much less missing values (Fig. 3a, center panel) and it more than doubled the number of available quantification data at low CV (Fig. 3a, bottom panel). It doubled sensitivity (Fig. 3b, top panel) and AUROC (Supplementary Fig. 5a, upper row) in DE analyses enabling detection of significantly more true positive spike-in proteins (Fig. 3b, center panel) while keeping false positive rates low (Fig. 3b, bottom panel). Spike-in ratios were comparably well estimated by MaxQuant and IceR in all tested scenarios with errors being at median always below 25% (Supplementary Fig. 5d). Generally, increasing the number of MS1 spectra by selecting a lower TopN method or by increasing gradient lengths is especially beneficial for IceR as it increases reproducibility of quantifications and sensitivity to detect differentially abundant proteins. IceR further improved sensitivity in case of very low sample injection amounts (Fig. 3b and supplementary Fig. 4e). Interestingly, median ion counts for quantified peptide features was highest in the 10 ng setup (Supplementary Fig. 5b), and peptide features that were quantified in all three injection amounts revealed that ion counts were only slightly lower in 10 ng than in 500 ng sample injections (Supplementary Fig. 5c). These collective data show that IceR delivers favourable results in a range of commonly used experimental regimens.

**Fig. 3.**
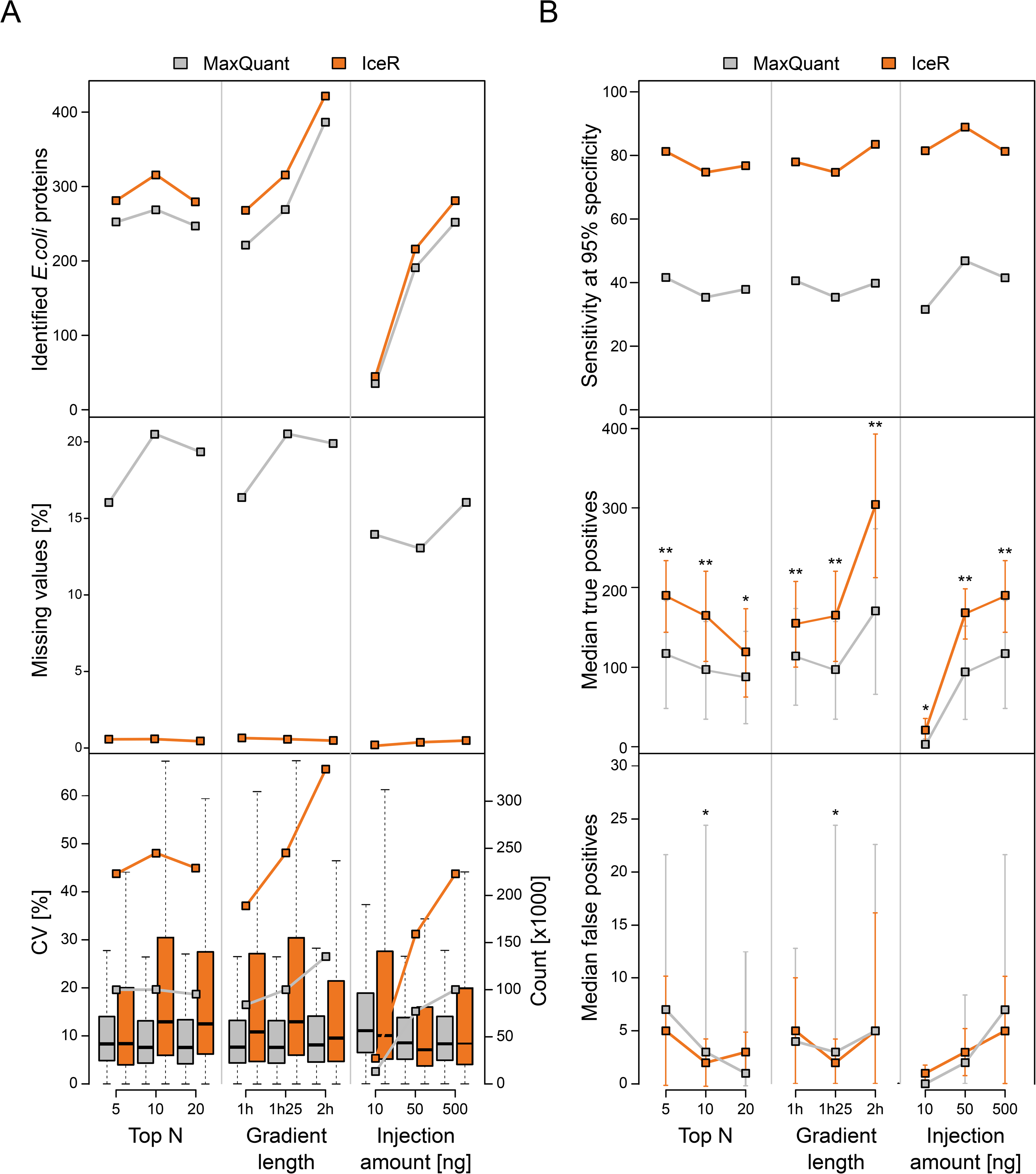
IceR enables robustly improved sensitivity for wide ranges of typically varied MS analysis parameters. **a**, Effect of TopN, gradient length and sample injection amount on numbers of identified *E.coli* proteins with at least two quantification events (top panel), missing values (center panel) and CVs of quantification (bottom panel) in MaxQuant (grey) and IceR (orange) data. Each condition was analysed in 3 replicates. **b**, as in **a** but showing effect of parameters on sensitivity (extracted from ROC curves at 95 % specificity, upper panel), detection of true positives (center panel) and detection of false positives (bottom panel) over all pairwise differential expression analyses in MaxQuant (grey) and IceR (orange) data. Asterisks indicate significance level: * p-value < 0.05, ** p-value < 0.01.

### Performance of IceR in comparison to DIA label-free proteomics

Data-independent acquisition (DIA) is an emerging technology for quantitative proteomics and has been shown to be superior in comparison to DDA as it can result in fewer missing values, and lower CVs across replicates^26^. Since IceR performed particularly well with regard to returning low missing value rates (Fig. 1c, Fig. 2b and Fig, 3a), we here wanted to evaluate the performance of IceR in comparison to DIA. To enable a more comprehensive comparison, we chose two data sets with different complexities: 1) a publicly available data set with 12 non-human proteins spiked into human lysate^27^, and 2) an in-house generated data set with a complete *E. coli* lysate spiked into human lysate.

The first data set generated by Bruderer et. al.^27^ was originally used to introduce a SWATH-MS-type DIA workflow called hyper reaction monitoring (HRM), and to compare its performance against DDA. For that purpose, 12 proteins were spiked into a constant human background introducing protein abundance changes ranging from as little as 10 % up to 5000 %, collectively resulting in 8 different samples (n=3) and 28 possible pairwise comparisons. All 12 spiked proteins were identified in the DDA and DIA data, and total numbers of identified proteins and peptides were comparable between MaxQuant, DIA (HRM) and IceR (Supplementary Fig. 6a,b). The fraction of missing values on peptide-level (19.5% in MaxQuant) could be reduced to 1.6% in DIA and, remarkably, to 0.6% in IceR data (Fig. 4a). Comparison of observed CVs on peptide-level between MaxQuant, DIA and IceR showed the highly reproducible quantification in DIA data. Median CV could be reduced from 16% in MaxQuant data to 8% in DIA data, along with an increase in available quantification events by 58% (Supplementary Fig. 6c). Strikingly, IceR resulted in a 250% increase in available quantification events compared to MaxQuant. Since this included more variable low abundant features, the median CV was slightly higher in IceR results (17.5%). When focusing on confidently quantified peptide features (pvalue of ion accumulation < 0.05, signal to background >= 4), median CV could be reduced to 10%, however, still with ~ 10% more quantification events compared to MaxQuant data. ROC curves revealed the highly improved performance of IceR (AUROC 0.90) compared to MaxQuant (AUROC 0.66) for detection of differentially abundant proteins (Fig. 4b), approaching the performance of DIA (AUROC 0.96). Of 336 total true positives (100%) over all 28 pairwise DE analyses (adj. pvalue < 0.01, absolute fold-change > 10%), IceR resulted in 64% cumulative true positives with significant differential abundance, compared to 46% for MaxQuant and 69% for DIA (Fig. 4c). DIA outperformed IceR especially due to its higher sensitivity for very small abundance differences (< 40%, Supplementary Fig. 6d). False positive rates were generally far below 1% for all methods (Fig 4c and supplementary Fig. 6e).

**Fig. 4.**
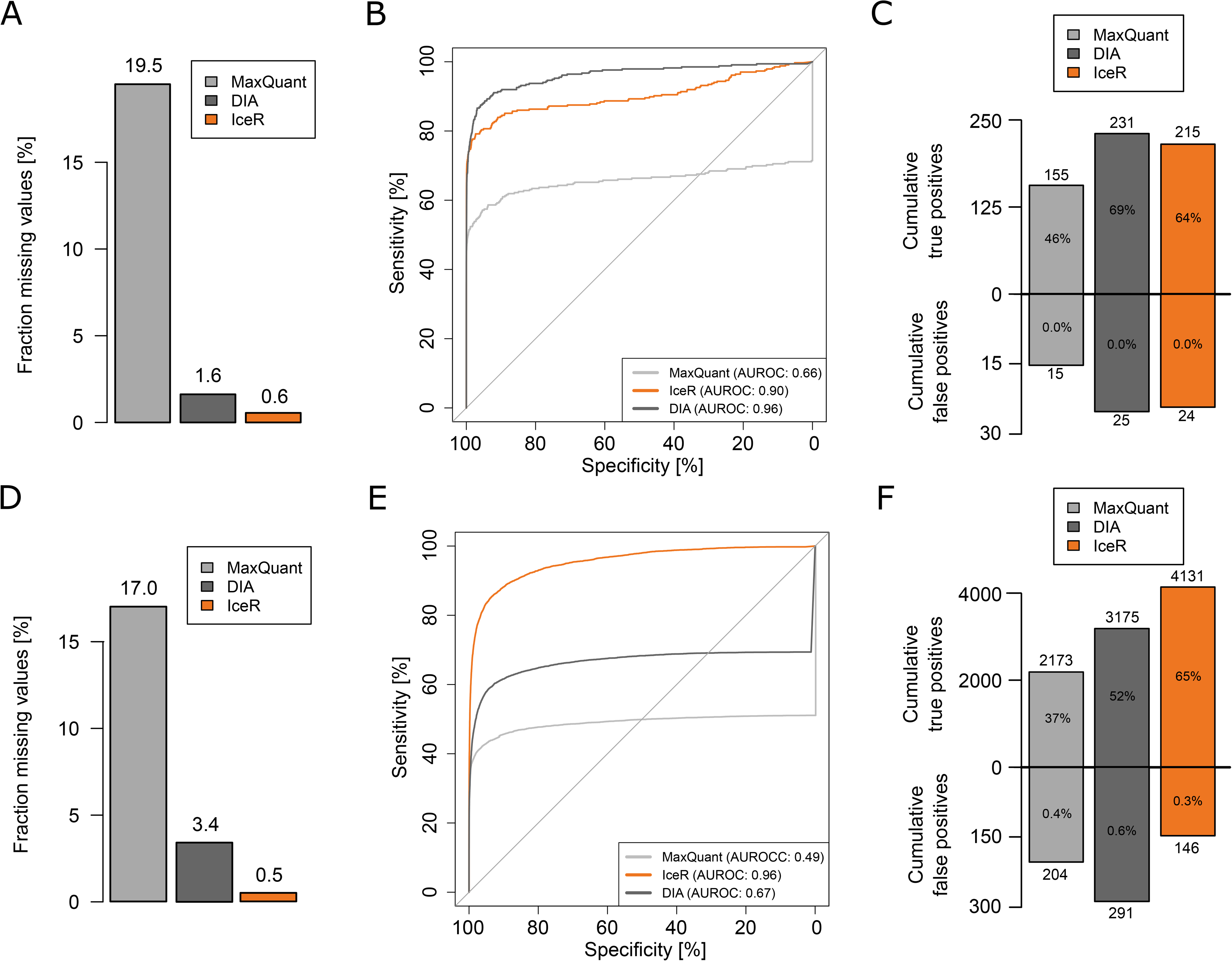
IceR enables label-free DDA proteomics with DIA performance. **a**, Fraction of missing values on peptide-level in MaxQuant (grey), DIA (red) and IceR (orange) results for a publicly available data set of 12 spiked proteins into a constant complex background at 8 concentrations (n=3). **b**, Receiver operating characteristic (ROC) over all pairwise differential expression analyses on protein-level in MaxQuant (grey) and on peptide-level in IceR (orange) and DIA (red) data. Corresponding areas under the ROC (AUROC) are indicated. **c**, Cumulative true and false positives over all (28) pairwise DE analyses for MaxQuant (grey), DIA (dark grey) and IceR (orange). True and false positive rates are indicated. **d**, Fraction of missing values on peptide-level in MaxQuant (grey), DIA (red) and IceR (orange) results for a data set of an *E. coli* lysate spiked into a constant background at 6 concentrations (n=3). **e**, Receiver operating characteristic (ROC) over all pairwise differential expression analyses protein-level in MaxQuant (grey) and on peptide-level in IceR (orange) and DIA (red) data. Corresponding areas under the ROC (AUROC) are indicated. **f**, Cumulative true and false positives over all (15) pairwise DE analyses for MaxQuant (grey), DIA (dark grey) and IceR (orange). True and false positive rates are indicated.

Next, we compared the performance of IceR, MaxQuant and DIA in the more complex situation of an *E. coli* lysate spiked into a human lysate at 6 different amounts as used above (0, 3, 4.5, 6, 7.5, and 9%). Samples were analysed in DDA and DIA mode with a 2-hour gradient method. The required spectral libraries for DIA were generated from DDA runs, and hence the total required MS resources for the DIA data set were increased (see methods for details). In total, about 400 *E. coli* proteins could be identified based on approximately 2800 peptides in both data sets (Supplementary Fig. 6f-g). The fraction of missing values was reduced from 17.0% in MaxQuant to 3.4% in DIA and even to 0.5% in IceR (Fig. 4d). Quantifications were most reproducible in DIA results, however, IceR resulted in many more available peptide quantification events (+165%) including more variable low-abundance peptide features (Supplementary Fig. 6h). Filtering IceR data for robust quantifications resulted in CVs comparable to DIA data. Pairwise DE analyses on peptide-level for DIA and IceR revealed the outstanding performance of IceR (AUROC 0.96) compared to MaxQuant (AUROC 0.49) and even to DIA results (AUROC 0.67) in this data set (Fig. 4e). Similarly, IceR resulted in the highest relative and absolute cumulative true positive rate over all DE analyses, greatly outperforming MaxQuant and DIA (Fig. 4f). The superior performance of IceR is driven by its capability to enable abundance ratio estimations even against low/no spike-in conditions and its improved sensitivity for small abundance ratios compared to MaxQuant (Supplementary Fig. 6i). In contrast, DIA fails to estimate ratios against no spike-in conditions but shows higher sensitivity for low abundance ratios (Supplementary Fig 6i). As before, false positive rates were generally far below 1% for all methods (Supplementary Fig. 6j).

In summary, IceR enables label-free DDA proteomic analyses with DIA-like performance with regard to sensitivity, absence of missing data, and quantitative accuracy. IceR consistently allows more sensitive detection of differential abundances in comparison to standard DDA data analysis and especially boosts sensitivity in case of comparisons against very low-abundance conditions.

### Addition of ion mobility dimension to label-free proteomics analyses

Combining IM separation with MS promises improved specificity and accuracy for PIP as recently demonstrated for the match-between-runs algorithm in MaxQuant^28^. Still, tools utilizing the additional ion mobility dimension for ion-based PIP are lacking. Here, we reanalysed our *E. coli* spike-in tool sample set on a timsTOF Pro mass spectrometer with a comparable 2-hour gradient method as described above. The total number of identified proteins could be increased by 30% compared to the Q Exactive HF (QE-HF) measurements while the total fraction of missing values was decreased from 17 % to 11 % on peptide-level in MaxQuant (4D match between runs enabled) (Fig. 5a). IceR reduced this to < 1 % missing values in both cases (Fig 5a). Over all pairwise DE analyses, MaxQuant identified 2212 (QE-HF) and 3931 (timsTOF Pro) true positive proteins, which represent 38 % and 51 % of the proteins with known differential abundance in these respective data sets (Fig. 5b). In contrast, IceR enabled detection of 4131 (65 % of total) and 6011 (78 % of total) true positives in QE-HF and timsTOF Pro data, respectively (Fig 5b). Interestingly, IceR even enabled the detection of more true positives in QE-HF data compared to MaxQuant in timsTOF Pro data. The superior sensitivity and specificity of IceR was also apparent by comparing receiver operating characteristics (ROC) and areas under the ROC (AUROC) for the respective conditions (Fig. 5c).

**Fig. 5.**
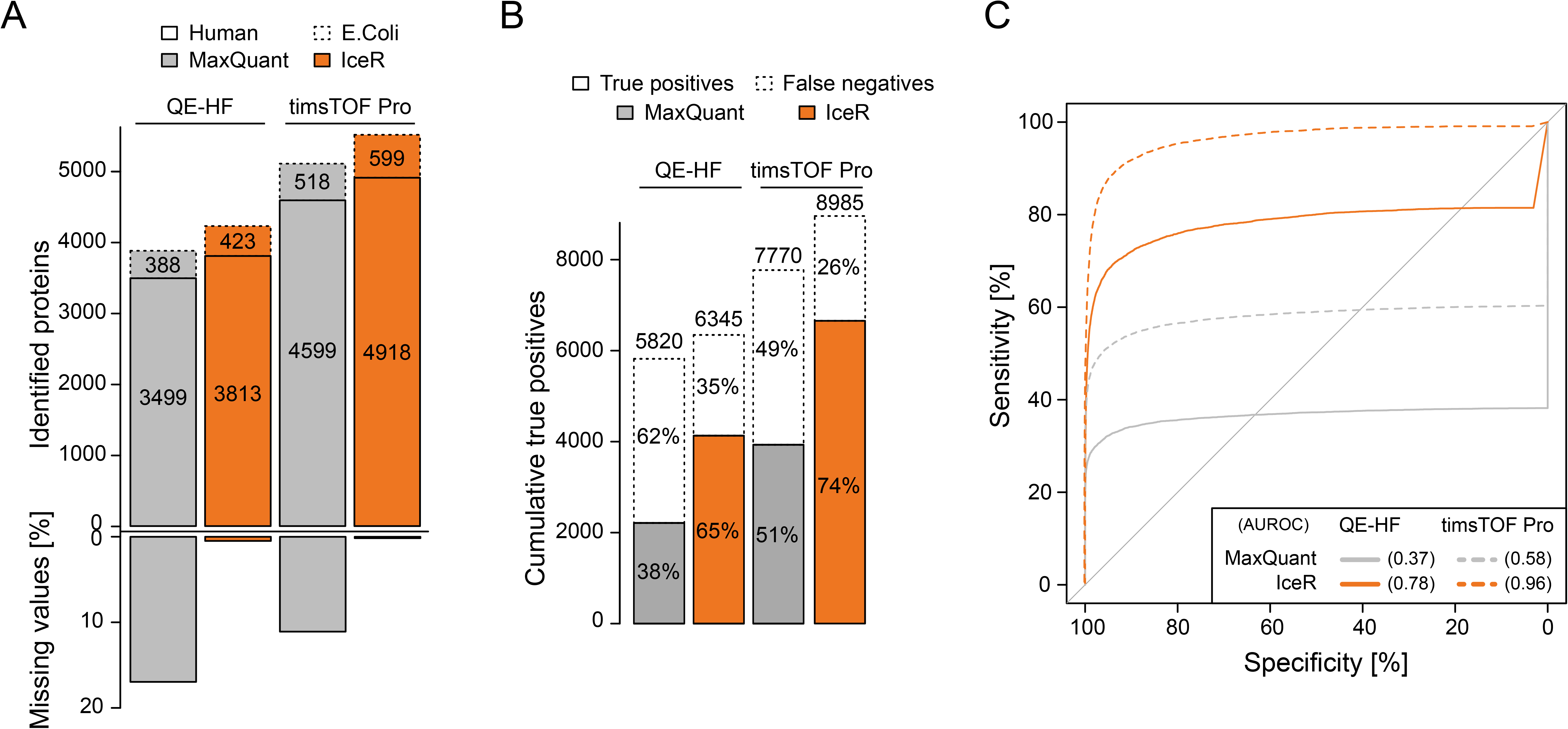
Application of IceR to timsTOF Pro proteomics data. **a**, Comparison of numbers of identified human (solid box) and *E. coli* (dashed box) proteins as well as fraction of missing values in the in-house generated *E. coli* spike-in data set analysed on a Q Exactive HF (QE-HF) and a timsTOF Pro followed by peptide and protein quantification using MaxQuant (grey) or IceR (orange). **b**, Number of detected differentially abundant (true positives, solid box) and missed (false negatives, dashed box) *E. coli* proteins cumulated over all pairwise differential expression analyses for data sets measured on a Q Exactive HF (QE-HF) or timsTOF Pro quantified by MaxQuant (grey), DIA (dark grey) or IceR (orange). Relative fractions from expected true differentially abundant proteins over all pairwise DE analyses per condition are indicated. In case of timsTOF Pro data, IceR was run with and without awareness of ion mobility (± IM) information. Cumulative false discovery rate per condition is indicated. **c**, Receiver operating characteristics (ROC) over all pairwise differential expression analyses for MaxQuant (light grey), DIA (dark grey) and IceR (orange) for data sets measured on a Q Exactive HF (QE-HF, solid lines) or timsTOF Pro (dashed lines). Area under the ROC (AUROC) per condition is indicated.

In summary, IceR highly benefits from the additional ion mobility dimension, lifting its sensitivity far above the MaxQuant workflow. Furthermore, here for the first time we showed the use of an ion-based PIP-implementing quantitative workflow on timsTOF Pro data.

### Application of IceR to plasma proteomics data

Blood plasma is an attractive and easily accessible body fluid for proteomics-based biomarker discovery, however, data completeness across samples tends to be low, due in part to the particular characteristic of the plasma proteome that the dynamic range of protein abundances is high. Here, we applied IceR to a publicly available plasma proteome data set of 32 finger prick samples acquired from one person over 8 consecutive days in short (20-min) LCMS analyses^29^. Originally, 257 proteins were quantified with at least 2 peptides by MaxQuant (MBR enabled) with a missing value rate of 11 % of the proteins per sample. In comparison, IceR enabled identification of 279 proteins where the fraction of missing values per sample could be reduced to at median 2 %. The highly improved data completeness from IceR enabled full quantification of 248 proteins (89 % of all identified proteins in IceR data) over all 32 samples while MaxQuant only allowed full quantification of 195 proteins (79 % of all identified proteins in MaxQuant data, Fig. 6a). The 53 additional proteins now rescued by IceR with 100% completeness showed reproducible quantification levels over all samples and ranged from low to high overall abundance (Fig. 6b) indicating that the IceR workflow goes far beyond simple missing-value imputation. These proteins included several important blood biomarkers like the indicators for myocardial infarction LDHA/LDHB, the coagulation factor F7, the liver injury marker CPS1, the infection marker CRP, the metabolic syndrome marker FABP5^30^ and the inflammatory predictor GSN^31^.

**Fig. 6.**
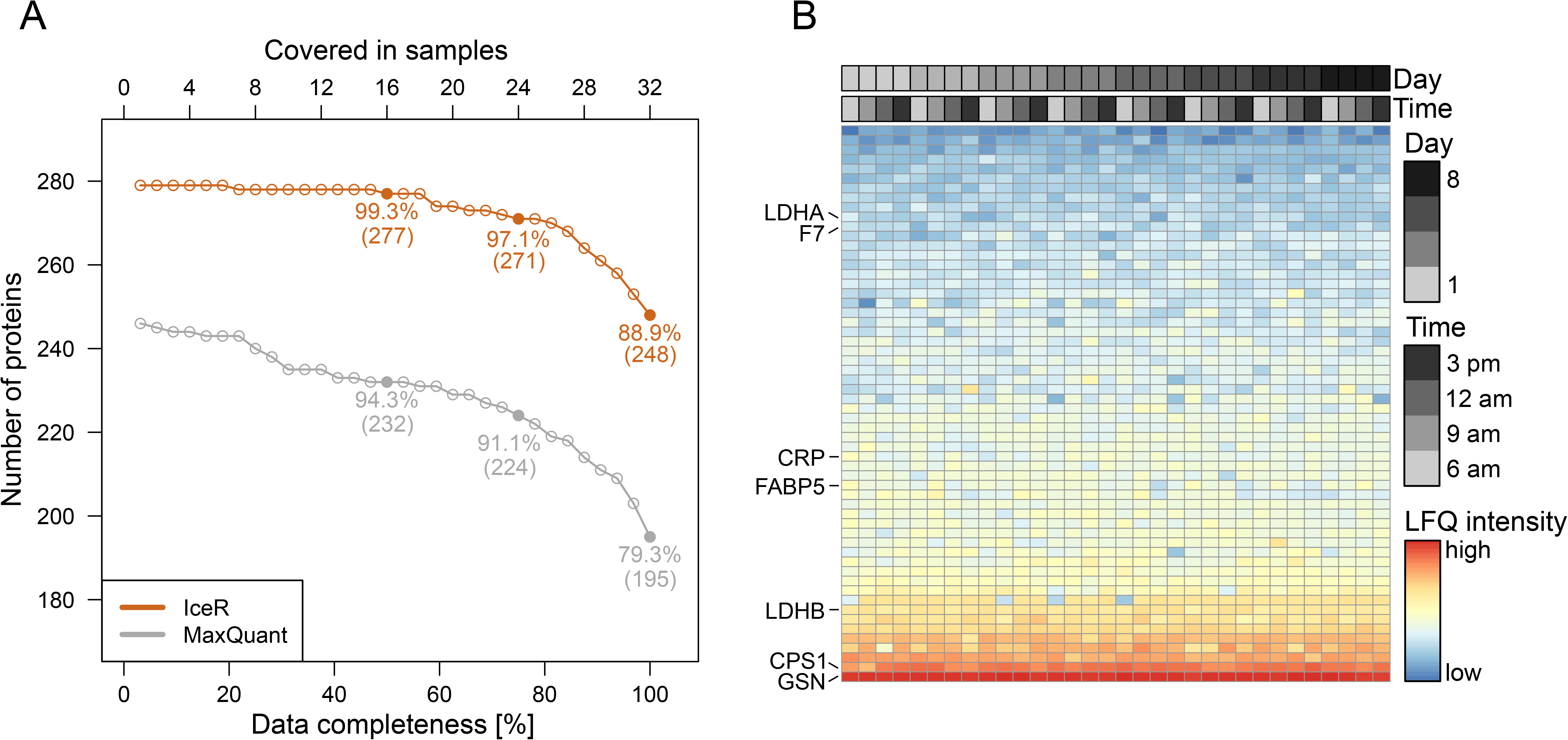
Application of IceR to a plasma proteomics data set. **a**, Numbers of fully quantified proteins with increasing number of samples for MaxQuant (grey) and IceR (orange) in plasma proteomics analyses of 32 finger prick samples acquired from one person over 8 consecutive days. Relative fractions of fully quantified proteins at 50, 75 and 100 % data completeness are highlighted. **b**, LFQ intensities of proteins without missing values in IceR but incomplete quantification in MaxQuant. Samples are ordered per sample acquisition day and time. Colors indicate protein abundance estimations. 7 important plasma protein markers are highlighted.

In summary, the improved sensitivity of IceR allows robust and reproducible quantification of more proteins with better data completeness in plasma proteome profiling data, thus displaying a more comprehensive portrait of a person’s health state.

### Application of IceR to label-free single-cell proteomics data

As demonstrated above, IceR allows increased sensitivity and data completeness even at very low sample injection amounts (Fig. 3 and supplementary Fig. 5). Since these are critical properties for low-input applications, we reasoned that IceR could be suitable to enhance sampling depth and data completeness of label-free single-cell proteomics data. To evaluate this, we selected a published data set from Zhu et al.^32^ where single developing hair and progenitor cells were isolated from utricles of embryonic chickens. Cells were separated by fluorescence-activated cell sorting (FACS) into FM1-43high hair cells and FM1-43low progenitor cells and subsequently processed using the nanoPOTS^33^ approach. In total, 10 single FM1-43high hair cells and 10 single FM1-43low progenitor cells were analysed in one batch. As two single hair and four single progenitor cell samples were previously determined to be empty^32^, these were here excluded from subsequent analyses. Along with the singlecell samples, pools of 20 FM1-43_high_ cells (n=3) and pools of 20 FM1-43_low_ cells (n=3) were used as matching libraries to boost identification rates in single-cell analyses. Nevertheless, and in line with the published data^32^, on average only 23 proteins (min: 5, max: 53) and 103 peptides (min: 35, max: 231) could be quantified per single-cell by MaxQuant (Fig. 7a). In contrast, when applying IceR to these data, on average 431 proteins (min: 417, max: 454) and 2651 features (min: 2511, max: 2769) could be quantified per single-cell sample (Fig. 7b and supplementary Fig. 7a). The critical contributor to this improvement is IceR’s distinct capability to rescue low-abundance features, thus decreasing the rate of missing values from 95.1% to 8.6% (protein-level) and from 95.3% to 3.6% (peptide-level), boosting both the number of identified/quantified peptides/proteins, and data completeness (Fig. 7c). Reproducibility of quantification was comparable between MaxQuant and IceR results, however IceR enabled in excess of 14-fold and 18-fold more quantification events on protein and peptide-level, respectively, compared to MaxQuant outputs (Supplementary Fig. 7b,c). t-distributed stochastic neighbour embedding (tSNE^34^) revealed clearer separation of hair and progenitor cells in IceR results (Silhouette score of 0.73) compared to MaxQuant outputs (Silhouette score of 0.53, Fig. 7d,e) demonstrating the improved data quality after requantification by IceR. This was even more obvious when performing DE analysis between single hair cells and single progenitor cells. While in MaxQuant data differential abundance could be tested only for 37 proteins due to the high data sparsity (Supplementary Fig. 7d), IceR enabled differential abundance testing for 491 proteins, and could detect increased expression of proteins known to be enriched in hair cells, including CALB2, CKB, and CRABP1 (Fig. 7f). Unsupervised *de novo* chronological ordering of cells using the R package CellTrails^35^ segregated all cells according to their FM1-43 uptake (Fig. 7g and Supplementary Fig. 7e). Further, developmental trajectories in MaxQuant and IceR results were used to examine protein expression dynamics as a function of developmental pseudotime (Fig. 7h and Supplementary Fig. 7f). Crucially, the highly improved data completeness after requantification by IceR allowed analysis of protein expression dynamics for 6-fold more proteins (413 vs 66), including proteins known to be enriched in progenitor cells (e.g. TPM3, AGR3, and TMSB4X) and in hair cells like (e.g. OTOF, CALB2, MYO6, CKB, AK1, CRABP1, and GAPDH) (Fig. 7i). Accumulated ion intensities in IceR-selected DICE windows followed the expected developmental trajectory, as observed for peptides originating from OTOF and TPM3 (Fig 6j) and several other proteins (Supplementary Fig. 7g,h), and clearly distinguished progenitor from hair cells. Enhanced sensitivity of protein detection enabled by IceR now allowed detection of additional proteins with systematic expression changes along the developmental pseudotime axis agreeing with previously published RNA expression^32^.

**Fig. 7.**
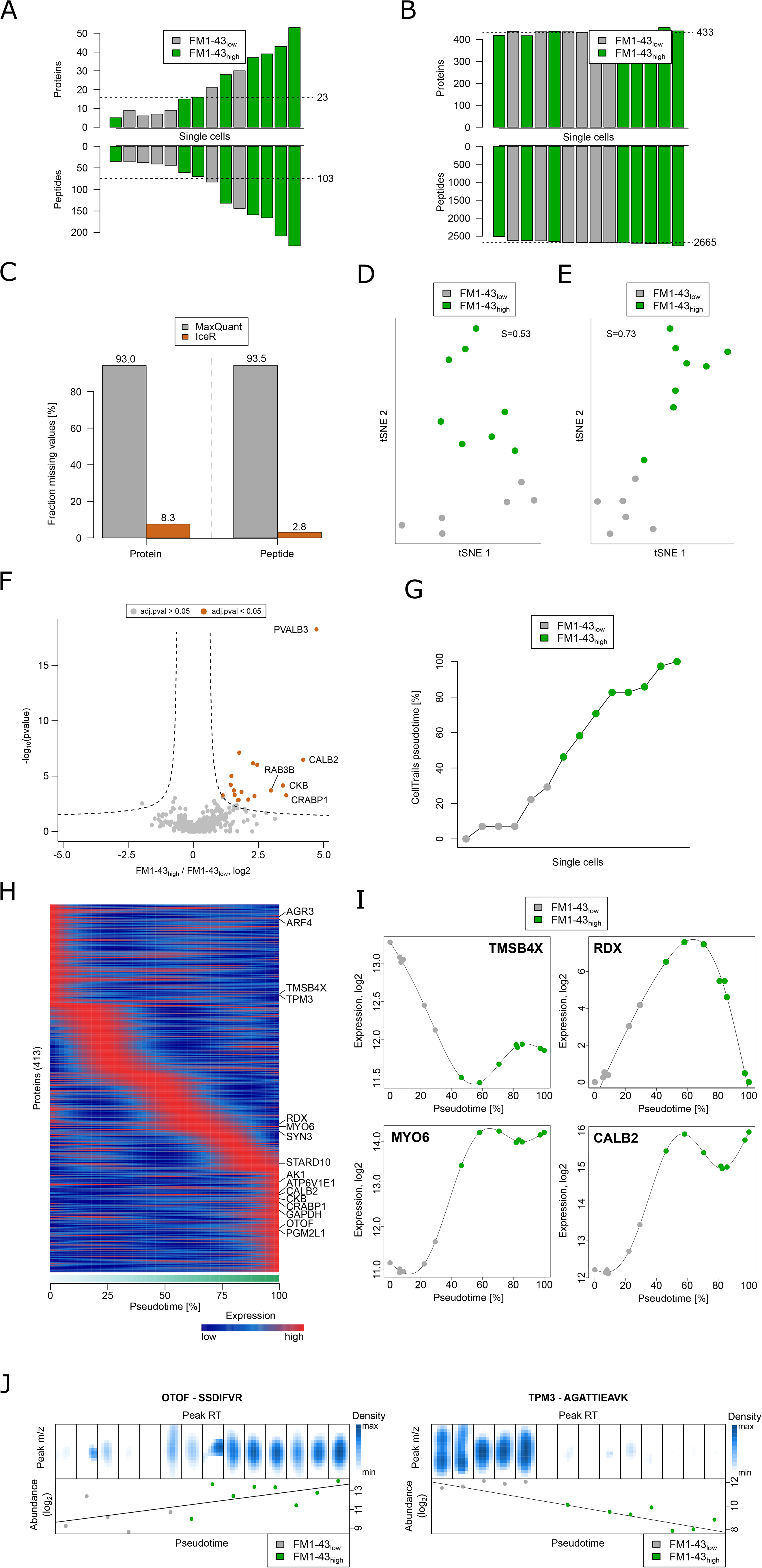
IceR boosts completeness and quality of single-cell proteomics data. **a**, Numbers of proteins (upper panel) and peptides (lower panel) and their average (dashed line) quantified by MaxQuant per single hair (FM1-43_high_, green) and progenitor (FM1-43_low_, grey) cell sample. **b**, as in a, but after data reprocessing by IceR. **c**, Fraction of missing values on protein- and peptide-level in MaxQuant (grey) and IceR (orange) results. **d**, Dimensional reduction of protein abundance estimations in MaxQuant data by t-distributed stochastic neighbour embedding (tSNE). Progenitor cells (FM1-43_low_) are coloured in grey. Hair cells (FM1-43_high_) are coloured in green. Silhouette score (S) is indicated. **e**, As in d but after data reprocessing by IceR. **f**, Volcano plot showing detected significantly (orange, adj. pvalue < 0.05, absolute fold-change > 2) differently abundant proteins between single hair cells and single progenitor cells in IceR data. Significance cutoff is indicated by a dashed line. **g**, Chronological ordering of single-cells as a function of CellTrails’ inferred pseudotime from IceR data. Single-cells are coloured according to their FM1-43 uptake. **h**, Scaled expression dynamics over pseudotime for all analysed proteins in IceR data based on generalized additive models (GAM). Low and high temporal protein expression is indicated by blue and red colour tones, respectively. **i**, Absolute expression dynamics of log2 expression levels as a function of pseudotime for various proteins. Single-cells are coloured according to their FM1-43 uptake. **j**, Ion density in IceR-selected DICE windows per single-cell sample (upper panels) for the peptides VLTLDLYK and AGATTIEAVK of OTOF (hair cell marker protein) and TPM3 (progenitor cell marker protein), respectively. Corresponding peptide abundances (lower panels) are ordered by CellTrails’ inferred pseudotime.

These overlaps included ARF4, SYN3, STARD10, ATP6V1E1, PGM2L1 and RDX (Fig. 7h). The latter is described to anchor cytoskeletal actin of stereocilia to hair cell membranes^36^. Interestingly, RDX showed highest transcript and protein expression midway through the developmental trajectory (Fig 6i), suggesting that a maturation-specific transient expression of this protein may be required for proper functionality. Importantly, with the exception of SYN3, none of the above-mentioned proteins could be revealed in the original paper either with regard to differential or pseudotime-dependent protein expression. This convincingly demonstrates that re-quantification by IceR uncovers biology that has remained hidden after analysis by conventional data analysis tools. In addition, it shows that DICE-based analysis via IceR is a promising strategy to boost performance of single-cell proteomics in general.

## Discussion

Our results show that IceR can greatly increase data completeness and quality in DDA-based quantitative label-free proteomics. Its greatly enhanced sensitivity and specificity could be demonstrated for a broad spectrum of published and newly generated data sets demonstrating its universal applicability. Although it had been previously shown that DICE-based data analysis has highly attractive performance features^1^, IceR is the first to offer this as a comprehensive yet userfriendly R-package. In addition, since DDA-based proteomics is widely used and firmly established, having undergone many rounds of methodological and hardware optimization over the last decade, and since IceR can be readily integrated in existing workflows, we expect that IceR can be easily adopted and that it will be of great value for the proteomics community. Furthermore, IceR improved the results obtained from a timsTOF Pro instrument, indicating that DICE-based analysis with IceR should be a valuable component in the emerging application of ion mobility mass spectrometry in proteomics.

We envision that IceR can have a strong impact in two main directions, that both receive considerable interest in present-day proteomics, namely biomarker discovery and low-input analysis. By necessity, biomarker discovery requires large sample cohorts, where, problematically, DDA-based proteomics returns decreasing numbers of fully quantified proteins with increasing cohort size^12^. We have shown here that IceR improved this situation in a series of plasma samples by simultaneously increasing proteome sampling depth and data completeness. Importantly, the property that IceR reduces missing value rates to those seen in DIA, and that it primarily rescues low-abundance proteins, demonstrates its potential to open up novel reservoirs of clinically relevant proteins in DDA data. In the area of low-input proteomics, the field enjoys considerable excitement with early examples demonstrating proof-of-principle of single-cell proteome analysis^16,33,37^. This is the combined result of miniaturized sample preparation, narrow-bore chromatography, and increased sensitivity in mass spectrometry, in conjunction with novel workflows such as TMT multiplexing with booster channels^38–40^. The fact that improved bioinformatics solutions have a major role to play in innovative single-cell strategies has remained underexposed, yet our data showed that re-analysis of an existing single-cell data set by IceR boosted the number of consistently quantified proteins from a few dozen to several hundreds. Indeed, this represents an improvement that has been difficult to achieve by any experimental innovation. As an upshot, highly increased sensitivity allowed the detection of many more differentially expressed proteins and allowed analyses of protein expression dynamics in greater detail. Importantly, this was done directly in single cells, thus indicating that label-free proteomic approaches can be a valuable alternative to TMT multiplexing approaches in the context of single-cell proteomics, avoiding shortcomings and controversies associated with the presence and magnitude of booster channels. In conclusion, IceR contributes to improved analysis of proteomic data in many experimental settings ranging from mainstream proteome characterization to biomarker discovery and low-input proteomics, where IceR could lift single-cell proteomics from infancy to early childhood.

## Acknowledgements

This work was supported by the Ministry of Education and Research (BMBF), as part of the National Research Node “Mass spectrometry in Systems Medicine” (MSCoreSys), under grant agreement 031L0212A.

## Author contributions

M.K. developed IceR; M.K. and T.M. designed experiments; T.M. performed experiments; M.K. analysed data; M.K. and J.K. wrote the manuscript; O.S. and J.K. supervised the work.

## Data availability

All data is available in the main text or the supplementary materials. The mass spectrometry proteomics data have been deposited to the ProteomeXchange Consortium via the PRIDE^41^ partner repository with the dataset identifier PXD019777 (username, password: xwk0ViB5). Protein sequences were taken from the UniProt database (https://www.uniprot.org/).

## Code availability

An implementation of the above described IceR procedure will be made available as an R-package at https://github.com/mathiaskalxdorf/IceR/

---

**Supplementary Fig. 1 – IceR workflow in detail.**

IceR starts from MaxQuant result files and MS-data in the mzXML format. If MS-data is supplied as Thermo Raw-files, they can be automatically converted to mzXML format by MSConvert from ProteoWizard. The IceR workflow consists of 13 steps, summarized here and described in more detail in Supplementary Information X.

Step 1) Alignment windows are determined by estimating deviations of observed retention time (RT) and calibrated m/z for peptides identified in samples.

Step 2) IceR features are defined by aggregating MaxQuant features across samples using defined alignment windows.

Step 3) Introduce decoy and +1-isotope features per IceR feature.

Step 4) RT- and m/z-correction factors are extracted per IceR feature and sample. If a feature was not detected by MaxQuant for an IceR feature in an individual sample, data modelling approaches are used to predict correction factors.

Step 5) Peptide sequence information within IceR features is propagated from sequenced MaxQ features between samples.

Step 6) Background noise, which is expected per IceR feature quantification, is estimated by counting and summing up intensities of ions that fall into decoy feature DICE-windows.

Step 7) Normal kernel density estimation to detect accumulations of ions (peaks) in RT- and m/z-space around the expected DICE-window per IceR feature and sample.

Step 8) Peaks are selected by applying a robust selection algorithm. Ions falling into selected DICE-windows are counted, distinguished into signal- or background-ions and corresponding intensities are summed.

Step 9) The significance of ion accumulation per quantification is determined by comparing the number of observed ions in DICE-windows against expected background noise ion count distributions (i.e. observed decoy feature ions). The quality of each quantification is further evaluated based on signal to noise ratios.

Step 10) Peak selection over samples is statistically evaluated and significant outliers are excluded.

Step 11) A peak selection false discovery rate (FDR) is estimated per sample.

Step 12) Optional: Missing quantifications of IceR features can be imputed using the decoy feature-based sample-specific background noise models.

Step 13) Peptide quantifications are aggregated to protein quantifications using the Top3, total sum intensity and MaxLFQ approach.

**Supplementary Fig. 2 – IceR quality control assessed by processing of iPRG 2015 data.**

**a**, Numbers of IceR features per sample sequenced by MSMS, by the match-between-runs algorithm of MaxQuant, by feature-based PIP or ion-based PIP by IceR. **b**, Density plot per sample visualizing accumulation of peptide ions (LWSAEIPNLYR) of the spiked protein lacZ detected by normal kernel density estimation. Black dashed boxes indicate expected 2D peak locations per sample. Detected 2D peaks are indicated with grey dashed boxes. True known peaks (by MS/MS identifications or PIP) in respective samples are indicated in green. These peaks are also selected by the peak selection algorithm of IceR for the respective sample. Peaks that were selected by the peak selection algorithm of IceR in samples lacking knowledge of the true peak location are indicate in orange. **c**, Deviation of IceR feature m/z and RT over samples before (raw) and after (aligned) alignment. **d**, False peak discovery rate estimation by masking location of 500 randomly picked IceR features with true identification per sample and evaluating how often a wrong peak is selected. Dashed line indicates mean false discovery rate (FDR). **e**, Numbers of monoisotopic (M+0, light colour) and +1-isotopic (M+1, dark colour) IceR features per sample quantified by DICE (orange), quantified by background noise model-based imputation (blue) or with missing values (grey). **f**, Fraction of missing values on peptide-level in DeMix-Q (red), MaxQuant (grey) and IceR (orange) outputs. **g**, Heatmap representation of quantified peptides of six spiked proteins in four tool samples (n=3) in MaxQuant (left), DeMix-Q (middle) and IceR (right) results. Peptides are ordered over all three outputs. Low abundance peptides are coloured blue, high abundant peptides are coloured red, and missing values are indicated in grey. **h**, Boxplot representation of log_2_ deviations between estimated abundance ratios and true spike-in ratios of respective pairwise differential expression analyses. Center line, median; box limits, upper and lower quartiles; whiskers, 1.5x interquartile range. **i**, Comparison of adjusted pvalues for spiked proteins in pairwise differential expression analyses in DeMix-Q and IceR outputs. Coefficient of determination (R^2^), slope and x-intercept of the linear model fit are indicated. *** indicates pvalue < 0.001.

**Supplementary Fig. 3 – Detailed IceR quality control assessment for iPRG 2015 data.**

**a**, Numbers of detected features with and without peptide sequence by MaxQuant per sample. **b**, Deviations in m/z and retention time between samples used to automatically define required feature alignment windows for IceR workflow (75%-quantile of m/z deviations and upper whisker of RT deviations). **c**, Smoothed scatterplot visualizing 20 % of observed feature m/z corrections over sample-specific random forest model predictions. Coefficient of determination (Rsq) is indicated. **d**, Smoothed scatterplot visualizing sample- and chromatographic retention time-specific generalized additive models (GAM) fitted to observed RT corrections per feature and sample. **e**, Smoothed scatterplot visualizing observed mean ion intensities within decoy ion direct ion current extraction windows over chromatographic retention time. Fitted GAM is indicate by a red line. Boxplot representations of mean ion intensities of decoy features, standard deviation of ion intensities of decoy features and numbers of ions per decoy feature are plotted. **f**, Boxplot visualization of determined significances of ion accumulations of IceR feature quantifications per sample. **g**, Boxplot visualization of signal to background ratios (log2) of IceR feature quantifications per sample. **h**, Absolute and relative fraction of IceR features per sample for which selected peaks are aligned to (orange) or are significantly deviating from (grey) corresponding peaks in all other samples. **i**, Absolute and relative fraction of +1-isotope IceR features for which peaks were selected with significant RT and/or m/z deviation compared to its respective monoisotopic feature. **j**, Number of proteins identified with at least 2 features by DeMix-Q (red), MaxQuant (grey) and IceR (orange). **k**, Number of peptides identified by DeMix-Q (red), MaxQuant (grey) and IceR (orange). **l**, Boxplot of coefficients of variation (CV) of peptide quantifications in DeMix-Q (red), MaxQuant (grey), IceR (orange), and filtered IceR (pvalue of ion accumulation < 0.01, signal to background >= 4, light orange) results. Numbers of available CVs per boxplot are indicated. Center line, median; box limits, upper and lower quartiles; whiskers, 1.5x interquartile range. **m**, True positive rates (TPR %) of pairwise differential expression analyses on protein- and peptide-level in MaxQuant (grey), DeMix-Q (red) and IceR (orange) data. **n**, As in m but showing corresponding false positive rates (FPR %).

**Supplementary Fig. 4 – Evaluation of IceR based on two publicly available tool data sets.**

Supplementary analysis results for the tool data set of Ramus et al. (a-f) and Shen et al. (g-l). **a**, Numbers of quantified proteins in MaxQuant (grey) and IceR (orange) results. **b**, Coefficient of variation of protein quantifications in MaxQuant (grey) and IceR (orange) results. Numbers of available CVs per boxplot are indicated. Center line, median; box limits, upper and lower quartiles; whiskers, 1.5x interquartile range. **c**, as in b but showing CVs of peptide quantifications. **d**, Deviation of determined from true protein abundance ratios of pairwise differential expression analyses on protein- and peptide-level in MaxQuant (grey), MaxQuant with imputation (dark grey) and IceR (orange) data. Numbers of spiked proteins for which abundance ratios could be determined per condition are indicated. **e**, Median true positive rates for MaxQuant (grey), MaxQuant with imputation (dark grey) and IceR (orange) in pairwise differential expression analyses binned by abundance ratio and reference spike-in sample. **f**, Median false positive rates for MaxQuant (grey), MaxQuant with imputation (dark grey) and IceR (orange) in pairwise differential expression analyses binned by abundance ratio and reference spike-in sample. **g**, Numbers of quantified proteins in MaxQuant (grey), IonStar (red) and IceR (orange) results. **h**, Boxplot of coefficients of variation (CV) of protein quantifications in MaxQuant (grey), IonStar (red) and IceR (orange) outputs. Numbers of available CVs per boxplot are indicated. Center line, median; box limits, upper and lower quartiles; whiskers, 1.5x interquartile range. **i**, as in h but showing CVs of peptide quantifications. **j**, Deviation of determined from true protein abundance ratios of pairwise differential expression analyses on protein- and peptide-level in MaxQuant (grey), MaxQuant with imputation (dark grey), IonStar (red) and IceR (orange) data. Numbers of spiked proteins for which abundance ratios could be determined per condition are indicated.

**Supplementary Fig. 5 – IceR enables robustly improved sensitivity for a wide range of typically varied MS analysis parameters.**

**a**, ROC curves for pairwise differential expression analyses in MaxQuant (grey) and IceR (orange) data at respective Top N (left, Top5 to Top20), gradient length (center, 1h to 2h) and sample injection amount (right, 10 ng to 500 ng). Upper panels show ROC curves with number of expected true positives set to the number of identified spiked proteins in MaxQuant data per setting. Lower panels show ROC curves with number of expected true positives set to the absolute maximum number of identified spiked proteins per experiment. Respective areas under the ROC (AUROC) are indicated. Respective sensitivities at 95 % specificity are indicated by squares. **b**, Boxplot representation of available data points per feature quantification in respective experiment and setting. Median counts are indicated. Center line, median; box limits, upper and lower quartiles; whiskers, 1.5x interquartile range. **c**, As in **b** but showing numbers of available data points per quantification of features detected over all sample injection amounts. **d**, Deviation of observed from true protein abundance ratio in pairwise differential expression analyses in MaxQuant (grey) and IceR (orange) outputs in respective experiment and setting. Absolute numbers and relative fractions compared to MaxQuant results of spiked proteins for which abundance ratios could be determined per condition are indicated.

**Supplementary Fig. 6 – Comparing performance of IceR and DIA for label-free proteomics.**

Supplementary analysis results for the tool data set of Bruderer et al. (a-e) and an in-house generated tool data set (f-j). **a**, Number of quantified proteins in MaxQuant (grey), DIA (red) and IceR (orange) data. **b**, Number of quantified peptides in MaxQuant (grey), DIA (red) and IceR (orange) data. **c**, Coefficient of variation of peptide quantifications in MaxQuant (grey), DIA (red), IceR (orange) and filtered IceR (pvalue of ion accumulation < 0.05, signal to background >= 4, dark orange) results. Numbers of available CVs per boxplot are indicated. Center line, median; box limits, upper and lower quartiles; whiskers, 1.5x interquartile range. **d**, Mean true positive rates for MaxQuant (grey, protein-level DE), DIA (red, peptide-level DE) and IceR (orange, peptide-level DE) outputs over true spike ratio of pairwise differential expression analyses. **e**, Mean false positive rates for MaxQuant (grey, protein-level DE), DIA (red, peptide-level DE) and IceR (orange, peptide-level DE) outputs over true spike ratio of pairwise differential expression analyses. **f**, Number of quantified proteins in MaxQuant (grey), DIA (red) and IceR (orange) data. **g**, Number of quantified peptides in MaxQuant (grey), DIA (red) and IceR (orange) data. **h**, Coefficient of variation of peptide quantifications in MaxQuant (grey), DIA (red), IceR (orange) and filtered IceR (pvalue of ion accumulation < 0.05, signal to background >= 4, dark orange) results. Numbers of available CVs per boxplot are indicated. Center line, median; box limits, upper and lower quartiles; whiskers, 1.5x interquartile range. **i**, Mean true positive rates for MaxQuant (grey, protein-level DE), DIA (red, peptide-level DE) and IceR (orange, peptide-level DE) outputs over true spike ratio of pairwise differential expression analyses. **j**, Mean false positive rates for MaxQuant (grey, protein-level DE), DIA (red, peptide-level DE) and IceR (orange, peptide-level DE) outputs over true spike ratio of pairwise differential expression analyses.

**Supplementary Fig. 7 – Application of IceR to a single-cell proteomics data set.**

**a**, Numbers of IceR features per single cell sample with significant (pvalue < 0.1) accumulation of ions. Dashed line indicates average over all samples. Single cell samples are coloured based on their FM1-43 uptake. **b**, Coefficient of variation of protein quantifications in MaxQuant (grey) and IceR (orange) data. Numbers of available CVs in respective data is indicated. **c**, Coefficient of variation of peptide quantifications in MaxQuant (grey) and IceR (orange) data. Numbers of available CVs in respective data is indicated. **d**, Volcano plot showing detected significantly (orange, adj. pvalue < 0.05) differently abundant proteins between single hair cells and single progenitor cells in MaxQuant data. Significance cut-off is indicated by a dashed line. **e**, Chronological ordering of single cells as a function of CellTrails’ inferred pseudotime from MaxQuant data. Single cells are coloured according to their FM1-43 uptake. **f**, Scaled expression dynamics over pseudotime for all analyzed proteins in MaxQuant data based on generalized additive models (GAM). Low and high temporal protein expression is indicated by blue and red colour tones, respectively. **g**, Ion density in IceR-selected DICE windows per single-cell sample (upper panels) for a peptide (VLTLDLYK) of CKB (hair cell marker protein) and corresponding peptide abundances (lower panel) ordered by CellTrails’ inferred pseudotime. Samples in which the peptide was identified by MaxQuant are indicated with an M in the corresponding upper panels. **h**, As in g but for a peptide (SDKPDMAEIEK) of TMSB4X (progenitor cell marker protein).

