## Supplementary material for "IceR improves proteome coverage and data completeness in global and single-cell proteomics": Materials and Methods

*Generation of an in-house tool spike-in data set*

HeLa cells were pelleted from cell culture, snap frozen in liquid nitrogen and stored at -80°C. Cell pellet was reconstituted in 100 μL of 0.1% RapiGest SF Surfactant (Waters) in 100 mM triethylammonium bicarbonate (TEAB, Sigma-Aldrich) and 1x protease inhibitor cocktail (PIC, cOmplete, Sigma-Aldrich). Each sample was probe-sonicated for 4x 15 seconds at 10% frequency with storage on-ice between cycles of homogenization (Branson Digital Sonifier). Samples were centrifuged at 15.000x g, 4°C for 30 minutes to pellet any remaining cell- and tissue-debris, followed by transfer of the supernatant into new reaction tubes and protein quantification using a bicinchoninic acid assay (BCA, Pierce – Thermo Scientific). Proteins were denatured for 5 minutes at 95°C. Disulfide-bonds were reduced with Dithiothreitol (DTT, 5 mM final concentration, Biomol) at 60°C for 30 minutes. Cysteine residues were alkylated using chloroacetamide (CAA, 15 mM final concentration, Sigma-Aldrich) at 23°C for 30 minutes. Reduced and alkylated proteins were digested overnight at 37°C in a table-top thermomixer at 500 rpm using sequencing-grade modified trypsin (Promega) in ddH_2_O. Upon overnight protein digestion, each sample was acidified to a final concentration of 1% trifluoroacetic acid (TFA, Biosolve Chimie) and incubated at 37°C and 500 rpm for 30 minutes, in order to cleave and precipitate RapiGest. Subsequently, samples were centrifuged at 15.000x g, at 23°C for 30 minutes to pellet the RapiGest precipitate and recover the peptide-containing supernatant to a new reaction tube. MS injection-ready samples were stored at -20°C. E. coli lyophilized sample (Bio-Rad) was re-suspended in ddH_2_O to achieve a stock concentrations of 2 μg/μL. 100 μL (200 μg) were incubated at 95°C for 5 minutes, followed by reduction and alkylation using Dithiothreitol (DTT, 10 mM final concentration) at 37°C for 1-hour and chloroacetamide (CAA, 40 mM final concentration) at 23°C for 45 minutes at 500 rpm. Reduced and alkylated proteins were digested overnight at 37°C in a table-top thermomixer at 700 rpm using sequencing-grade modified trypsin (Promega) in ddH_2_O. Upon overnight protein digestion, each sample was acidified to a final concentration of 1% trifluoroacetic acid (TFA, Biosolve Chimie). MS injection-ready samples were stored at -20°C. Spike-in samples were prepared by mixing HeLa sample with 0%, 3%, 4.5%, 6%, 7.5% or 9% (wt/wt) of E. coli sample (n=3).

For the QE-HF acquisition, peptides were separated using the Easy NanoLC 1200 fitted with a trapping (Acclaim PepMap C18, 5μm, 100Å, 100 μm x 2cm, Thermo Fisher Scientific) and an analytical column (nanoEase MZ BEH C18, 1.7 μm, 130 Å, 75 μm x 25 cm, Waters). The outlet of the analytical column was coupled directly to a Q-Exactive HF Orbitrap (Thermo Fisher Scientific) mass spectrometer. Solvent A was ddH_2_O (Biosolve Chimie), 0.1% (v/v) FA (Biosolve Chimie) and solvent B was 80% acetonitrile (ACN, Pierce – Thermo Scientific) in ddH_2_O, 0.1% (v/v) FA. The samples were loaded with a constant flow of solvent A at a maximum pressure of 800 bar, onto the trapping column. Peptides were eluted via the analytical column at a constant flow of 0.3 μL/minute at 55°C using three different methods as described below. 2-hours method: During the elution, the percentage of solvent B was increased in a linear fashion from 3 to 8% in 4 minutes, then from 8% to 10% in 2 minutes, then from 10% to 32% in a further 68 minutes, and then to 50% B in 12 minutes. Finally, the gradient was finished with 8 minutes at 100% solvent B, followed by 11 minutes 97% solvent A. 1-hour 25 minutes method: During elution, the percentage of solvent B was increased linearly from 4 to 5% in 1 minute, then from 5% to 27% in 30 minutes, and then from 27% to 44% in a further 5 minutes. Finally, the gradient was finished with 10.1 minutes at 95% solvent B, followed by 13.5 minutes at 96% solvent A. 1-hour 10 minutes method: During elution, the percentage of solvent B was increased linearly from 3 to 8% in 4 minute, then from 8% to 10% in 2 minutes, and then from 10% to 32% in a further 17 minutes, and then to 50% B in 3 minutes. Finally, the gradient was finished with 8 minutes at 100% solvent B, followed by 11 minutes at 97% solvent A. Peptides were introduced into the mass spectrometer via a Pico-Tip Emitter 360 μm OD x 20 μm ID; 10 μm tip (New Objective) and at a spray voltage of 2 kV. The capillary temperature was set at 275°C. Full scan MS spectra with mass range m/z 350 to 1500 were acquired in the Orbitrap with a resolution of 60,000 FWHM. The filling time was set to a maximum of 50 ms with an automatic gain control (AGC) target of 3 x 10^6^ ions. The top 5, top 10, or top 20 most abundant ions per full scan were selected for an MS^2^ acquisition. Isotopes, unassigned charges, and charges of +1 or >+8 were excluded. The dynamic exclusion list was with a maximum retention period of 15 seconds (1-hour 10 minutes) or 25 seconds and a mass tolerance of plus and minus 10 ppm. For MS^2^ scans, the resolution was set to 15,000 FWHM with automatic gain control of 1 x 10^5^ ions and maximum fill time of 50 ms. The isolation window was set to 2 Th, with a fixed first mass of m/z 110, and stepped collision energy of 26.

For the timsTOF Pro PASEF acquisition of E. coli spike-in samples, peptides were separated using the Bruker nanoElute system fitted with an analytical column (Aurora Series Emitter Column with CSI fitting, C18, 1.6 μm, 75 μm x 25 cm) (Ion Optics). The outlet of the analytical column with a captive spray fitting was directly coupled to a timsTOF Pro (Bruker) mass spectrometer using a captive spray source. Solvent A was ddH2O (Biosolve Chimie), 0.1% (v/v) FA (Biosolve Chimie), 2% acetonitrile (ACN) (Pierce, Thermo Scientific), and solvent B was 100% ACN in ddH2O, 0.1% (v/v) FA. The samples were loaded at a constant maximum pressure of 900 bar. Peptides were eluted via the analytical column at a constant flow of 0.4 μL per minute at 55°C. During the elution, the percentage of solvent B was increased in a linear fashion from 2 to 17% in 60 minutes, then from 17 to 25% in 30 minutes, then from 35 to 37% in a further 10 minutes, and then to 95% in 10 minutes. Finally, the gradient was finished with 10 minutes at 95% solvent B. Peptides were introduced into the mass spectrometer via the standard Bruker captive spray source at default settings. The glass capillary was operated at 3500 V with 500 V end plate offset and 3 L/minute dry gas at 180°C. Full scan MS spectra with mass range m/z 100 to 1700 and a 1/k0 range from 0.6 to 1.6 V*s/cm2 with 100 ms ramp time were acquired with a rolling average switched on (10x). The duty cycle was locked at 100% and the TIMS mode was enabled. All timsTOF and nanoElute methods were default provided by Bruker.

*Data repositories*

Raw LC-MS/MS data of the individual publicly available data sets were downloaded from respective sources:

1. iPRG2015 – <ftp://iprg_study:ABRF329@ftp.peptideatlas.org/>
2. Ramus et al. – <http://proteomecentral.proteomexchange.org/cgi/GetDataset?ID=PXD001819>
3. Shen et al. – <http://proteomecentral.proteomexchange.org/cgi/GetDataset?ID=PXD003881>
4. Bruderer et al. – <ftp://PASS00589:WF6554orn@ftp.peptideatlas.org/>
5. Zhu et al. – <http://proteomecentral.proteomexchange.org/cgi/GetDataset?ID=PXD014256>
6. Geyer et al. - <http://proteomecentral.proteomexchange.org/cgi/GetDataset?ID=PXD002854>

The in-house generated LC-MS/MS raw files have been deposited to the ProteomeXchange Consortium via the PRIDE partner repository (Perez-Riverol et al., 2019) with the dataset identifier PXD019777.

*Data preprocessing*

Raw files were processed using MaxQuant (version 1.5.1.2 or 1.6.14.0). The searches for the individual data sets were performed against the following databases concatenated with reversed sequences:

1.) iPRG2015 – supplied iPRG2015 database (6634 entries)
2.) Ramus et al. – UniProt database consisting of reviewed S. cerevisiae and human UPS1 proteins (August 2019, 9805 entries)
3.) Shen et al. – UniProt database consisting of reviewed human and *E. coli* proteins (September 2019, 24644 entries)
4.) Bruderer et al. – UniProt database consisting of reviewed human proteins (August 2017, 20214 entries) including the 12 spiked proteins
5.) Zhu et al. – UniProt database consisting of chicken proteins (April 2020, 34726 entries)
6.) Geyer et al. - UniProt database consisting of reviewed human proteins (March 2020, 20365 entries)
7.) In-house generated *E. coli* spike-in data set – UniProt database consisting of reviewed human and *E. coli* proteins (September 2019, 24644 entries)

The Andromeda search engine was used with the following search criteria: enzyme was set to trypsin/P with up to 2 missed cleavages. Carbamidomethylation (C) was selected as a fixed modification; oxidation (M), acetylation (protein N-term) were set as variable modification. Match between runs was enabled with match time window set to 1 min and alignment time window set to 20 min. The search type for protein quantification was set to standard. Quantification intensities were calculated by the default fast MaxLFQ algorithm with minimal ratio count set to 1 or 2. Require MS/MS for LFQ comparisons was disabled. Peptide and protein hits were filtered at a false discovery rate of 1%, with a minimal peptide length of 7 amino acids. Second peptide search for the identification of chimeric MS2 spectra was enabled. Not mentioned MaxQuant settings were left as default.

*IceR workflow*

IceR requires raw files to be converted into centroided mzXML files. This can either be done by the user beforehand or IceR triggers the conversion using the ProteoWizard tool msConvert if installed. The complete workflow of IceR is illustrated in Supplementary Fig. 1 and includes 13 steps which are implemented in a R-package:

1. Features detected by MaxQuant are aligned over samples. For that purpose, deviations of observed retention time (RT) and calibrated m/z for peptides identified in samples are determined. The RT feature alignment window is defined as 1.5 x inter-quartile range of observed absolute RT deviations of peptides between samples. The m/z feature alignment window is defined as the inter-quartile range of observed absolute m/z deviations of peptides between samples. If a minimal RT-window or m/z-window is defined by the user and the observed window is smaller than the specified window, the user-defined minimal window is used. If the final RT-window and/or m/z-window is already specified by the user, the user-defined parameter is used.
2. Features detected by MaxQuant (MaxQ features from allpeptides.txt) are aggregated using the determined RT- and m/z-alignment windows. First, MaxQ features with same peptide sequence, same charge-state and same PTM over samples are aggregated and a new IceR feature with median RT and m/z of these MaxQ features is defined. Peptide features from samples with m/z or RT deviating from these medians by more than the defined alignment windows are excluded. Detected unknown MaxQ features within the alignment windows but without sequencing information are added. RT peak widths per IceR feature are defined as the maximum RT peak width observed for any of the aggregated MaxQ features. Overlapping IceR features (default: delta mass < 0.002 Da) are merged. Optional: Unknown features (without sequence information) which are left after peptide feature aggregation can be aggregated accordingly.
3. For every IceR feature, a decoy feature is generated. Here, alignment windows multiplied by 5 are added to the IceR feature RT and m/z. By default: For every IceR feature an expected +1-isotope feature is added. Here it is assumed that the isotope features should show an m/z shift of roughly 1.002 Da per charge.
4. For every IceR feature and every sample the individual m/z correction factor is extracted. For samples without an observed MaxQ feature in the respective IceR feature, the m/z correction factor has to be estimated. For that purpose, random forest models (RFs, R-package randomForest, version 4.6.14) are trained per sample based on 80 % of available MaxQ features with RT, m/z, charge and resolution as predictors and deviation of uncalibrated to calibrated m/z as response factors. Number of trees is set to 100. Number of variables randomly sampled as candidates at each split is set to 4. Minimal size of terminal nodes is set to 100. Trained models are validated using remaining 20 % of available data. Next, for every IceR feature and every sample the individual RT deviation of observed MaxQ features from IceR RT (median) are extracted. For samples without an observed MaxQ feature in the respective IceR feature the RT deviation has to be estimated. For that purpose, RT-dependent generalized additive models (GAMs, R-package mgcv, version 1.8.31) are fitted per sample to deviations between IceR feature RT and observed MaxQ feature RTs.
5. Peptide sequence information within IceR features is propagated (feature-based PIP) from sequenced MaxQ features between samples.
6. Background noise, which is expected per IceR feature quantification, is estimated by counting and summing up intensities of ions which are falling into decoy feature direct ion-current extraction (DICE)-windows. These windows are defined as m/z of IceR feature +/- m/z alignment window and RT +/- RT peak width / 2. Finally, RT-dependent GAMs are fitted to the observed decoy feature intensities and decoy ion counts are used to estimate number of ions which are randomly falling into DICE-windows (background noise).
7. Accumulations of ions (peaks) in RT- and m/z-space around the expected DICE-window per IceR feature and sample are detected by normal kernel density estimation (KDE, function kde2d in R-package MASS, version 7.3-51.5) and subsequent 2D-peak detection (local maxima). By default, KDE is performed with a resolution (grid points per dimension) of 50 and locations of up to 5 peaks (sorted by distance from expected peak location) with at least n detected ions (n = median decoy ion count) or localized within expected DICE-window are stored. Increasing KDE resolution improves the resolving power to detect peak locations, however, comes at the cost of longer processing times (doubling the resolution in theory results in quadruplicated processing times).
8. For every IceR feature, a peak per sample is selected. For samples with detected MaxQ feature, the peak closest to the expected peak location is selected as long as at least n ions form the peak (n = median decoy ion count) and the peak is located within the expected DICE-window. These peaks are called known. In all other cases, the peak closest to known peaks (in other samples), which is not overlapping with any other peak observed in samples with known peaks, is selected as long as its m/z is not deviating more than 3 times the m/z alignment window and its RT is not deviating more than the RT alignment window. If no peak is fulfilling these criteria for a sample, the expected DICE-window for this IceR feature is selected. Finally, all ions within the selected DICE-windows are counted and their intensities are summed. This total intensity is further distinguished into signal and background ion intensities by defining ions with an intensity higher than the background noise at respective RT (background noise GAM) + 2x standard deviation of background noise (decoy feature quantifications) as signal ions.
9. The significance of ion accumulation per quantification is determined by comparing number of observed ions in DICE-windows against expected background noise ion count distributions (observed decoy feature ions). All IceR features per sample with e.g. a quantification pvalue < 0.05 show a significantly higher accumulation of ions than expected by chance and are thus regarded as truly present. The quality of each quantification is further evaluated based on signal to noise ratios.
10. Peak selections are controlled and outliers removed. Two filters are applied: 1.) Features showing significantly increased interquartile ranges for peak RT or peak m/z between samples are completely removed. 2.) Features showing significantly deviating peak RT or m/z in individual samples are excluded. An additional filter is applied for +1-isotope IceR features by detecting outliers which show a significant deviation of peak RT or m/z between the monoisotopic and +1-isotope IceR features.
11. For every IceR analysis, its peak selection accuracy is estimated by performing a false discovery rate (FDR) analysis. For that purpose, 500 sequenced IceR features are randomly selected per sample, their known true peak locations are masked (treated as if no MaxQuant feature was detected) and it is then evaluated, how often the algorithm ends up selecting a wrong peak with deviating intensity.
12. Optional: IceR feature quantifications without any ion falling into the DICE-window result in missing values e.g. in case of true absence of a peptide. In this case the decoy feature-based sample-specific background noise models can be used to impute missing values with feature-specific background noise intensities. By default, data is imputed, however, none-imputed data is processed in parallel.
13. Peptide quantifications are aggregated to protein quantifications. The user can decide between the Top3, total sum intensity and MaxLFQ approach.

IceR results are stored as tab-delimited text files.

*Data of additional quantitative workflows*

To enable comparison of IceR against DeMix-Q and IonStar at their optimal parameter settings for the respective tool data sets, published quantitative workflow results were used:

- DeMix-Q – iPRG2015 data set - <https://www.mcponline.org/lookup/suppl/doi:10.1074/mcp.O115.055475/-/DC1/mcp.O115.055475-1.xlsx>
- IonStar – Shen et al. data set - <https://www.pnas.org/highwire/filestream/808767/field_highwire_adjunct_files/2/pnas.1800541115.sd02.xlsx>

Similarly, for comparison of IceR against HRM-DIA published by Bruderer et al., quantitative workflow results were downloaded from: https://www.mcponline.org/highwire/filestream/35126/field_highwire_adjunct_files/2/mcp.M114.044305-3.xlsx

*Data filtering and normalization*

The following filtering criteria were applied to respective quantitative workflow outputs for data sets 1, 2, 4 and 6:

- Remove contaminants, reverse hits, proteins identified only by PTM peptide and proteins identified with less than 2 peptides
- Keep protein quantifications per sample based on at least 2 peptides/features.

In case of data set 3 the respective quantitative workflow outputs the following filtering criteria were applied:

- Remove contaminants, reverse hits, proteins identified only by PTM peptide and proteins identified with less than 2 unique peptides
- Keep protein quantifications per sample based on at least 2 unique peptides.

In case of data set 5 the respective quantitative workflow outputs the following filtering criteria were applied:

- Remove contaminants, reverse hits, proteins identified only by PTM peptide and proteins identified with less than 2 (1 in case of DE and CellTrail analyses on MaxQuant data) peptides
- Keep protein quantifications per sample based on at least 2 (1 in case of DE and CellTrail analyses on MaxQuant data) peptides/features.

Subsequent to data filtering, protein and peptide quantities of samples of respective quantitative workflow outputs were median-normalized. In case of spike-in data sets, normalization factors were calculated based on constant background proteins.

*General data analysis*

Data analyses were performed and results were visualized using R (Version 3.6.3). For differential expression analyses on protein level, a modified t-test (R-package limma, version 3.42.2) was applied. Differential expression analyses on peptide level were performed using peptide-level expression-change averaging (R-package PECA^1^, version 1.22.0, ordinary t-test, ratio based on Top5 abundant features). Receiver operating characteristics (ROC) and areas under the ROC (AUROC) were utilized to compare performances of quantitative workflows for detecting differentially abundant proteins (R-package pROC, version 1.16.2).

*Single cell analysis*

Missing values of protein quantifications in MaxQuant data were imputed by random draws from a Gaussian distribution centered to the 1%-quantile of observed values per sample (R-package imputeLCMD, version 2.0). Dimensional reduction for visualization was performed using t-distributed stochastic neighbor embedding (tSNE, R-package tsne, version 0.1.3). Clustering of single cells was evaluated using the Silhouette score (R-package cluster, version 2.1.0). Unsupervised de novo chronological ordering of cells was performed using the R-package CellTrails (29874578, version 1.4.0).

*Code availability*

The proteomics quantification workflow IceR will be made available on GitHub (<https://mathiaskalxdorf.github.io/IceR/>).

*References*

1.	Suomi, T., Corthals, G. L., Nevalainen, O. S. & Elo, L. L. Using Peptide-Level Proteomics Data for Detecting Differentially Expressed Proteins. *J. Proteome Res.* **14**, 4564–4570 (2015).
