## Supplemental Text for "IceR improves proteome coverage and data completeness in global and single-cell proteomics"

*IceR analysis pipeline*

Below, we explain the consecutive steps performed by IceR in more detail, accompanying Suppl Fig 1 showing these steps in a graphically. In addition to describing the analysis workflow, performance of IceR is illustrated based on its processing of a published spike-in data set (iPRG 2015^1^). In that study, yeast digests were spiked (n=3) with different concentrations of six exogenous marker proteins. Its limited complexity typically allows good performances of analysis workflows to detect spiked proteins. As this data set was previously used to evaluate DeMix-Q^2^, it enabled us to directly compare results of DeMix-Q and IceR. MaxQuant results of the same data were used as a reference.

The IceR approach includes the following 13 steps, aligning with the panel numbers in Suppl. Fig 1:

1.) Estimation of feature alignment windows. In the iPRG2015 data set, peptide sequences could be assigned to only 20% of all features detected by MaxQuant, thereby leaving 80% of features unused (Supplementary Fig. 3a). IceR estimated suitable RT- and m/z-feature alignment windows by determining deviations in RT and m/z for identified peptide features over samples. Here, the m/z- and RT-alignment windows were determined to be at least 0.35 mDa and 0.35 min, respectively (Supplementary Fig. 3b).
2.) MaxQuant features with same peptide sequence, charge-state, and PTM over samples are aggregated into a new IceR feature with median RT and m/z. Peptide features from samples with m/z or RT deviating from these medians by more than the defined alignment windows are excluded.
3.) For an increased statistical power and quantification accuracy, for every IceR feature an expected +1-isotope feature is added. Furthermore, for estimation of background noise per quantification, a decoy feature with an arbitrarily shifted extraction window is generated for every IceR target feature.
4.) The original location (RT and m/z and if available IM) for every IceR feature in every sample is determined, and its distance to the aggregate IceR feature locations is stored as a correction factor. If MaxQuant feature information is lacking, this location has to be estimated. For that purpose, m/z- and RT-corrections are modelled by sample-specific random forests (RF) and generalized additive models (GAM), respectively (Supplementary Fig. 3c,d). The majority of iPRG2015 samples show consistent chromatographic elution profiles while e.g. sample 7 shows an increasing shift in retention times which would prevent PIP.
5.) Unsequenced MaxQuant features that fall into the alignment windows of an IceR feature are regarded as missed peptide features. For these, sequence information is transferred based on feature-based PIP. In case of the iPRG2015 data set, sequence information could be transferred by feature-base PIP for about 34% of IceR features lacking an identity in the iPRG2015 data set (Supplementary Figure 2a). For all remaining IceR features per sample (on average 13%) that even lack detection of any MaxQuant feature, sequence information could be transferred by ion-based PIP.
6.) Decoy features are used to model and estimate quantification background noise by counting and summing up intensities of ions that randomly fall into DICE-windows (Supplementary Fig. 3e).
7.) Locations of up to 5 peaks (accumulations of ions) around the expected DICE-windows of IceR features (sorted by distance from the expected peak location) are detected by kernel density estimations and subsequent peak detection. Kernel density plots and detected 2D peaks for a peptide (LWSAEIPNLYR) of the spiked protein lacZ are shown in Supplementary Fig. 2b. MaxQuant (match-between-runs enabled) detected this peptide only in samples 4 – 9 while IceR was able to detect the correct peak in all 12 samples after its 2‑step alignment procedure (Supplementary Fig. 2b).
8.) IceR tries to selects a peak for every aggregated feature in all samples. In case of IM data, available ions are prefiltered requiring their IM to fall within the expected IM window. For samples with detected MaxQuant feature, the peak closest to the expected feature location is selected as long as it is formed by n ions (n = median decoy ion count) and is located within the expected DICE‑window. These peaks are classified as ‘known’. In all other cases, IceR picks the peak closest to all known locations in other samples as long as it is not overlapping with any other peak, its m/z is not deviating more than 3 times the m/z alignment window and its RT is not deviating more than the RT alignment window (Supplementary Fig. 2b). If no peak is fulfilling these criteria for a sample, the expected DICE-window for this IceR feature is selected. The improved alignment approach of IceR reduced the median deviations of RT and m/z between samples by more than 3-fold and 2-fold, respectively (Supplementary Fig. 2c). After peak selection, all ions within the selected DICE-windows are counted and their intensities are summed.
9.) The significance of ion accumulation per quantification is determined by comparing the number of observed ions in DICE-windows against background noise ion count distributions. Resulting quantification pvalues thus indicate if more ions than expected by chance accumulated within respective DICE-windows. Features with significant accumulations (pvalue<0.05) can be regarded as truly present. The quality of each quantification is further evaluated based on signal to noise ratios. For the current data set, almost all quantifications show significant ion accumulation (median –log_10_ pvalues of 5, Supplementary Fig. 3f) and a generally good signal to noise ratio (median S/N ~ 14, Supplementary Fig. 3g). +1-isotope IceR features that show no significant accumulation of ions (pvalue>0.05) in any sample are removed.
10.) Peak selections are verified and outliers removed. Two filters are applied: 1.) Features showing significantly (pvalue<0.05) increased interquartile ranges for peak RT or peak m/z between samples are completely removed. These numbers are usually below 1% (Supplementary Fig. 3h) 2.) Features showing significantly (pvalue<0.05) deviating peak RT or m/z in individual samples are excluded. An additional filter is applied for +1-isotope IceR features by detecting outliers that show a significant (pvalue<0.05) deviation of peak RT or m/z between the monoisotopic and +1-isotope IceR features (typically below 20% (Supplementary Fig. 3i)).
11.) For every IceR analysis, accuracy of peak selection is estimated by performing a false discovery rate (FDR) analysis. Therefore, 500 IceR features with peaks classified as ‘known’ are randomly selected per sample, their true peak locations are masked (treated as if no MaxQuant feature was detected) and it is then evaluated, how often the algorithm ends up selecting a wrong peak with deviating intensity. Typically, peak selection FDR is below 1%. In case of the iPRG2015 data set, an average FDR of 0.6% was estimated (Supplementary Fig. 2d).
12.) Optionally, the background noise models can be used to impute missing feature quantifications. Missing values that resulted from the applied filtering criteria are not imputed because of likely wrong peak selection. In the iPRG2015 data set, on average 98% of monoisotopic peptide features could be directly quantified by DICE and remaining 2% of quantifications were imputed (Supplementary Figure 2e). In contrast, MaxQuant could estimate abundances only for 86% of all identified peptides. Additionally added +1-isotope features by IceR could be directly quantified by DICE in 84% of cases while 2% of quantifications were imputed. The remaining 14% of +1-isotope IceR features are left with missing values due to exclusion by filtering criteria.
13.) Finally, protein-level quantification is obtained by aggregating feature level quantifications. By default, aggregation is performed by Top-3^3^, total sum intensity and MaxLFQ^4^ approaches.

Following IceR requantification, we compared the results of our workflow with the outputs generated by MaxQuant (match-between-runs enabled) as well as with published DeMix-Q results. Numbers of identified proteins and peptides were comparable between all three approaches (Supplementary Fig. 3j,k). However, MaxQuant resulted in 12-fold more missing values compared to DeMix-Q and IceR (Supplementary Fig. 2f,g). DeMix-Q showed lowest coefficients of variation (CVs) of peptide quantifications, however, IceR resulted in almost 2-fold more available quantification events including more variable low abundant features (Supplementary Fig. 3l). When focusing on feature quantifications with significant ion accumulation and signal-to-background ratio (quantification pvalue < 0.01, S/B > 4‑fold), CVs in IceR were comparable to DeMix-Q results. To take advantage of the highly increased numbers of peptide quantifications available in DeMix-Q and IceR results, differential expression (DE) analyses were performed on peptide-level using peptide-level expression-change averaging (PECA, 26380941) for all tools. Additionally, as peptide-level DE can result in reduced sensitivity in case of high data sparsity as typically returned from standard label-free quantification workflows, DE analyses were also performed on protein-level for MaxQuant and IceR. The limited complexity of the data set enabled similar performances for detecting true positives by all three tools (Supplementary Fig. 3m,n). However, IceR resulted in most accurate and precise protein abundance ratio estimations (Supplementary Fig. 2h) and enhanced statistical power for DE analyses compared to DeMix-Q (Supplementary Fig. 2i).
